# A new approach for high-content traction force microscopy to characterize large cell ensembles

**DOI:** 10.1101/2025.07.04.663196

**Authors:** Nicolas Desjardins-Lecavalier, Santiago Costantino

**Affiliations:** Maisonneuve-Rosemont Hospital Research Center, 5415, boulevard de l’Assomption, Montréal, QC H1T 2M4, Canada; Institut de Génie Biomédical, Université of Montréal, Pavillon Paul-G.-Desmarais, 2960, chemin de la Tour, Montréal, QC H3T 1J4, Canada; Department of Ophthalmology, University of Montreal, Pavillon Roger-Gaudry, Bureau S-700, 2900, boul. Édouard-Montpetit, Montréal, QC H3T 1J4, Canada

## Abstract

1.1

Accurate measurements of cellular forces are important for understanding a wide range of biological processes where traction plays a major role. The characterization of mechanical properties is needed to unravel complex phenomena like migration, morphogenesis, mechanotransduction, or shape regulation, but accurate data on large numbers of single cells remain scarce and challenging. The capacity to measure forces in populations of cells and to identify subsets within heterogeneous ensembles would enable to reveal and manipulate their intrinsic complexity. Traction Force Microscopy (TFM) is a technology that can quantify the contractile forces exerted by cells via measuring the displacement of fluorescent beads embedded on the surface of a soft substrate with precisely defined mechanical properties. However, conventional numerical approaches for measuring cellular forces using TFM are labor-intensive and can yield significant artifacts, making them ill-suited for high-throughput analysis. In this work, we propose using the Demons algorithm instead, leading to significant improvements in both computational efficiency and accuracy. Based computer simulations, we show that in some situations this methodology outperforms conventional approaches in terms of speed, it is less sensitive to the blur induced by out-of-focus images and improves the accuracy of force field reconstructions. Additionally, we conducted experiments using cell lines and gels of distinct stiffness to demonstrate that the Demons algorithm is capable of real-time analysis and is effective at clustering cells according to their mechanotype.

**Statement of significance:** Large cellular ensembles encompass a wide variety of traits that require detailed but high-throughput technologies for characterisation. Granular descriptions of cell populations enable to describe their heterogeneity are essential to understand biological phenomena in the search of effective treatments. Most advances in the field of Traction Force Microscopy (TFM) are oriented towards high resolution force measurements at sub-micron adhesion scale that reveal the intricacies of molecular interactions. Here, we propose a strategy to exploit TFM for characterizing the mechanotypes of large cell ensembles. We benchmark and show the capacities of using the Demons algorithm to measure bead displacement fields, the improvement in force reconstructions obtained, and experimentally demonstrate that single cell analysis of cell dynamics can be used to cluster and describe different cell types.

## 1.3 Introduction

Measurement of traction is required to study the ability of cells to sense, generate and transmit forces to their microenvironment and neighboring cells. Cells initiate intracellular signaling cascades in response to external mechanical stimuli that impact their fate and functions, such as adhesion dynamics (1–4), cell shape (5), proliferation, differentiation and metabolism (6, 7). On a macroscopic scale, coordinated changes in cell shape lead to force propagation throughout tissues, which is essential to morphogenesis; these forces can also serve as cues for immune response and wound healing (8–10). On a microscopic scale, migrating single cells contract, stretch and apply forces as they maneuver through complex environments. These shape transitions are influenced by expression of cytoskeletal and adhesion proteins, and the mechanical properties of the substrate. In the case of disease, migrating metastatic cells represent an important determinant of progression: they navigate through densely packed microenvironments, exerting pressure to penetrate the extracellular matrix, distort, and relocate to invade distant organs (11). Taken together, the thorough characterization of cellular traction is important to understand the role of mechanical forces on biological phenomena that determine both organismal homeostasis and the outbreak of maladies.

Most biological ensembles display highly heterogeneous traits, thereby requiring single-cell characterization of many cells for a complete description. High-content force measurement techniques are needed to describe the wide variety of mechanical phenotypes that are found experimentally. Notably, even cells from a genetically homogeneous background display epigenetic and expression variance that yield substantial changes in force phenotypes; cancer cells are even more heterogenous (12), due to elevated mutation rates and chromosomic alterations (13, 14). High-content techniques must characterize not only the details of single adhesions or individual cells but assess the variability within cell populations and the capacity to probe critical but rare events.

The study of the signaling pathways, molecular properties and factors that determine the mechanotype has been an active research field for decades (15–17), highlighting key molecular players involved in cellular force generation. Recently, researchers have been trying to dissect the mechanical heterogeneity found in metastatic cell populations by isolating cells based on various properties like their adhesiveness (18), speed (19), or migration ability through a transwell (20).

Traction Force Microscopy (TFM) is the most widely used technique for quantifying cellular contractility, relying on readily available reagents and tools to measure the force fields generated by cells. In TFM, fluorescent beads are embedded on the surface of a soft hydrogel prior to cell seeding. As cells adhere and exert forces through contraction and extension, these forces cause bead displacements, which can be translated into traction forces using the known mechanical properties of the hydrogel.

Although a few studies have explored high-throughput TFM approaches—typically sacrificing spatial resolution by using low-magnification objectives (21, 22) —TFM is predominantly employed for high-precision measurements on a limited number of cells. This is due to the reliance on laser scanning microscopy and the computational intensity of bead displacement analysis. Recent advancements in TFM have further emphasized high-resolution, sub-adhesion scale force measurements, leveraging nanoscopy techniques(23).

It is widely recognized that errors in the bead displacements can propagate and lead to important artifacts in the force reconstruction (24), thus, a common numerical approach to minimize errors is to perform a parametrized Thikonov regularization in the force calculation. However, inaccuracies cannot be fully corrected with regularization and the precise assessment of the bead displacement field is still of prime importance. Computationally intensive cross-correlation techniques are typically used to measure the bead displacements but are slow to calculate and may result in significant error rates in force reconstruction, often underestimating traction by approximately 50% (25). Recently, Optical Flow techniques, which are faster to compute, have been introduced to effectively measure small displacements, and demonstrated superior performance compared to cross-correlation methods for this specific application (26).

The Demons algorithm, which we analyze here, is novel in TFM, but has been extensively utilized in the domain of medical image processing (27, 28). Demons iteratively traces spatio-temporal gradients of intensity to converge on a displacement field between two consecutive images and uses regularization by applying a Gaussian filter at each iteration, leading to a diffusion-like behavior. Here, we benchmark the Demons algorithm and demonstrate that it can be used to increase the throughput of TFM and minimize bead displacement errors (27, 28). We show, using simulated data, that this approach outperforms both the gold standard cross-correlation techniques and Optical Flow. Finally, we implement high content TFM to highlight the capacity to characterize the mechanoptype of two different cancer cell lines.

## 1.4 Methods

### 1.4.1 Computing displacement fields

In this study, we compare three computational techniques for measuring bead displacements and evaluating the precision of the resulting force field reconstructions. While cross-correlation methods are widely used, they are computationally intensive. In contrast, Optical Flow and the Demons algorithm offer significantly faster computation times, enabling high-throughput, real-time analysis of displacement fields, as illustrated in Figure S1. The Demons algorithm requires approximately five times fewer operations per pixel than cross-correlation (based on a 16×16 pixel template and linear sampling every 4 pixels, with 10 iterations), and nearly 6 times more than Optical Flow (with 5 iterations). Detailed estimations of floating-point operations are provided in the Supporting Materials section titled *Estimation of the Number of Floating-Point Operations*.

### 1.4.2 Cross-correlation

Cross-correlation methods have been widely employed in TFM, with numerous variations developed over time. Cross-correlation methods seek to localize regions of similarity between a moving image and a reference fixed image within a specified neighborhood; they are effectively used to determine the optimal displacement compatible with the deformation observed when beads are pulled. A reference template window ***W***_*ref,ci*_*(m*_*1*_,*m*_*2*_) is extracted from the reference fixed image at coordinates [*m*_*1*_,*m*_*2*_]. A second window, ***W***_*ci*_*(m*_*1*_*+b*_*1*_, *m*_*2*_*+b*_*2*_), is then extracted from the moving image, where coordinates are translated from [*m*_*1*_,*m*_*2*_] by [*b*_*1*_,*b*_*2*_].

The cross-correlation matrix ***cc***(*b_1_, b_2_*) is computed by assessing the correlation between ***W***_*ref,ci*_*(m*_*1*_,*m*_*2*_) and ***W***_*ci*_*(m*_*1*_*+b*_*1*_,*m*_*2*_*+b*_*2*_) across the defined neighborhood. This is mathematically represented as:

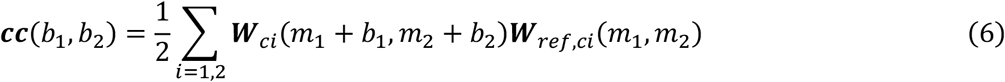

Sub-pixel displacements 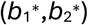 are then determined by fitting a Gaussian to the cross-correlation matrix and locating its maximum (29, 30). In this work, we employed a pyramidal implementation of Particle Image Velocimetry (PIV), specifically adapted for TFM applications (31). For all experiments, we used interrogation windows of 32, 16, and 8 pixels for the first, second, and third pyramid levels, respectively. Optimization of these parameters is discussed in detail in Section 1.7.

### 1.4.3 Optical Flow

Optical Flow refers to the distribution of velocities associated with the movement of brightness patterns within a scene as perceived by an observer (32). The method assumes that a moving object maintains a constant brightness and that displacement is small. Optical Flow constraint is expressed as (33):

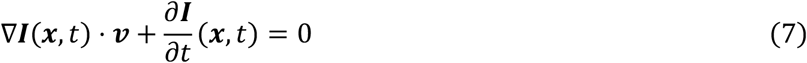

where *∇* ***I*** represents the spatial intensity gradient of the image, ***v*** is the velocity of the Optical Flow, and ***I***_*t*_ is the partial derivative of the intensity with respect to time. Consequently, the component of the Optical Flow in the direction of the brightness gradient (*∇* ***I***) can be formulated as:

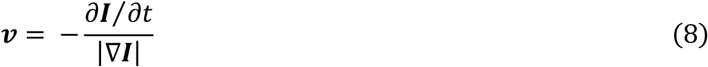

A wide range of motion estimation techniques are based on the concept of Optical Flow (34). In this study, we employed the Kanade-Lucas-Tomasi (KLT) variant (26, 35), implemented in MATLAB’s Vision Toolbox, which is commonly used within the TFM research community (26). The displacement ***v*** is determined by minimizing the following expression over a small spatial neighborhood *Ω*:

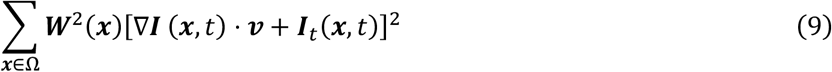

where ***W***(***x***) denotes a square function that regularizes *v*.

### 1.4.4 The Demons algorithm

Demons is a non-rigid image registration technique inspired by Maxwell’s demon thought experiment (27). To determine the pixel displacements that registers a moving image to a reference image, this approach iteratively calculates and applies a series of small transformations. In each iteration a Gaussian smoothing is applied to the transformation, mimicking a diffusion process (27, 28). While the technique performs the best in measuring displacements of freely diffusing particles, it also applies to beads embedded in an elastic medium (36). The small transformation, caused by “demonic” forces, makes the pixels move along the temporal and spatial gradients of the images, following the equation:

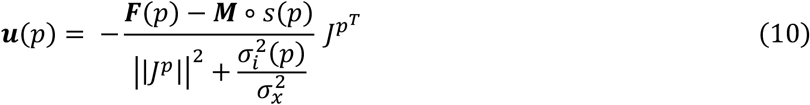

where ***u*** represents the iterative small displacement, ***F*** is the reference fixed image, ***M*** is the moving image, and *s* denotes the cumulative transformation up to the current iteration. The term *J*^*p*^ refers to the Jacobian, while *σ*_*i*_*(p)=*|***F****(p)-****M****°s(p)*| quantifies the noise in the image. Additionally, the constraint ||***u****(p)*||≤ *σ*_*x*_ accounts for the spatial uncertainty associated with the transformation. The smoothing parameter, which represents the standard deviation of the Gaussian, can be tuned. In this work, we used the MATLAB function imregDemons with only one pyramid level, a total gaussian smoothing of 0.75 pixels and 100 iterations. These values were obtained after numerical testing and optimization.

### 1.5 Reconstruction of traction forces

The bead displacements between the reference image (without cells) and the moving image (gel under contractile activity of cells) enable the reconstruction of the force field. We solved the inverse problem of the Boussinesq approximation to the elastostatic equation in the Fourier domain, which allows fast computation (24, 37). This approach limits the calculations to a 2-dimensional plane but greatly accelerates the computation of the traction forces.

Briefly, the formulation of the stress reconstruction works as follows: the strain tensor inside an infinite linearly elastic material can be written in the form of a 3×3 matrix of elements ϵ_*i,j*_ that is a function of the local displacement ***u*** = [*u*_1_, *u*_2_, *u*_3_] and spatial coordinates ***x*** = [*x*_1_, *x*_2_, *x*_3_]:

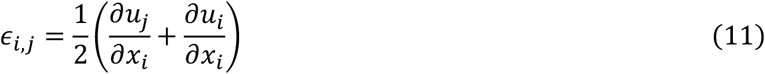

where *i* and *j* are the indices of the 3 dimensions.

For linearly elastic materials, the stress *σ*_*ij*_ is proportional to the strain *ε*_*ij*_.

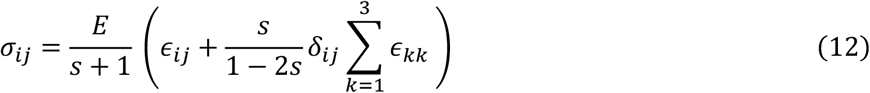

where *E* is the Young’s modulus, *s* is the Poisson ratio and *δ*_*ij*_ the Kronecker delta.

The Boussinesq approximation to the elastostatic equation (37) assumes a semi-infinite medium with stresses applied at the surface, which simplifies the problem, as forces and displacements in 2D only can be expressed as:

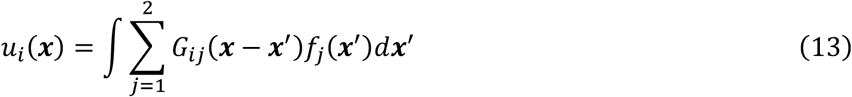

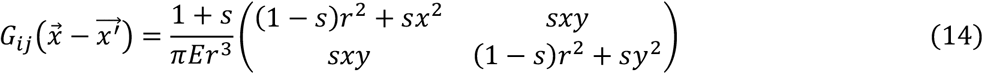

where *r =* |***x***|, *s* is the Poisson ratio, *G* the Green’s tensor for the Boussinesq approximation. In this work, we considered a Poisson ratio of 0.45 for polyacrylamide gels.

In the Fourier domain we can easily solve for the force (25, 38):

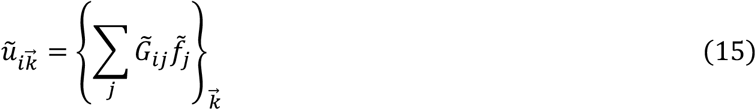

Where:

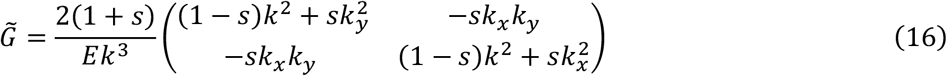

Where *k = (kx, ky)* is the wave number. To obtain the force field, the inverse problem must be solved. Importantly, this is an ill-posed problem, and even minor discrepancies in displacement data can significantly alter the resulting force estimation. Regularization approaches are used in the force field calculation to obtain smooth solutions despite the noise in the displacement field measurement (24, 39, 40). Thikonov regularization, where force magnitude is penalized, is typically used by requiring the tuning of a regularization parameter *L*, where we minimize:

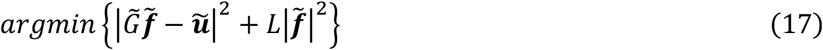

We typically solve numerically the problem under the following form (41):

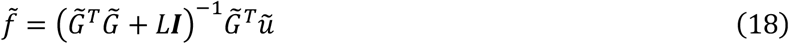

Where ***I*** is the identity matrix. For automatic selection of the parameter, a Bayesian approach has been developed which requires a statistical measurement of displacement noise performed outside the cell, where the displacements are assumed to be null (39).

## 1.6 Cellular traction simulations and error definitions

To compare the various methods used to measure bead displacements, we simulated bead images in two different scenarios: one where all beads are homogeneously displaced in the same direction, and one where we simulated a realistic displacement field generated by a distribution of focal cell adhesions.

### 1.6.1 Homogeneous displacement field

Beads where displaced homogeneously from 0 to 20 pixels with a step of 0.25 pixels. To mimic well-resolved vs. blurred beads, we adjusted the standard deviation of a Gaussian point spread function (PSF) to 0.33 pixels, 1.81 pixels and 10 pixels. We also varied the bead density, ranging from sparse 0.01 px^−2^, to 0.5 px^−2^, to a dense 1 px^−2^. For all the 84 displacement values, 3 PSF sizes and 3 bead densities, we generated 100 images by randomly shuffling the bead location. For each method, we fine-tuned and optimized parameters, i.e. the widow size for each pyramid level for cross-correlation, the block size for Optical Flow, and the total smoothing for Demons, and the and kept them constant throughout the experiments, independently from the size of the displacements, the PSF or the bead density. Figures S3-S11 show the evolutions of bead displacement RMSE, DTM and DTA for the algorithm parameters. We selected the parameters that represented the best compromise between the RMSE and the DTA. One displacement error measure was extracted per image, comparing the simulated ground truth to the measured displacement field. The error of the measurement was expressed using the root mean square error (RMSE). Detailed mathematical description of the error follows in subsection 4.3.3.

### 1.6.2 Cell-like traction field

To evaluate the impact of the accuracy of bead displacement measurements with each technique on the force reconstruction, we simulated the displacement field that is typically observed for a distribution of focal adhesions (FA) of a polarized cell. We segmented focal adhesions (FA) from a microscopic image of migrating breast cancer cells, specifically MDA-MB-231 cells (Figure 2a), and simulated equal forces for every adhesion pointing towards the center of mass of the cell (Figure 2b). We used the Boussinesq approximation to determine the motion of each bead (Figure 2c) and varied the traction, ranging it between 200 to 2000 Pascal, which resulted in average particle displacements under the FAs of 1-10 pixels, respectively. Similarly to the previous experiment, we replicated the experiment for 4 PSF standard deviation to values (0.33, 1, 2 and 5 pixels), and 4 bead densities (0.01, 0.1, 0.5 and 1 pixels^−2^). For each traction value, PSF size and bead density, we generated 200 images.

We then measured bead displacement errors and force reconstruction errors for each image by comparing the simulated to the reconstructed traction fields, and the simulated and measured bead displacement fields.

We selected

### 1.6.3 Error measurements

To quantify the bead displacement errors, we measured a root mean square error (RMSE) per image

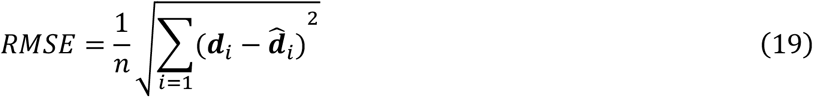

Where 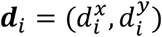 and 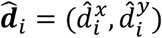 represent the simulated and measured displacements for the i^th^ pixel and *n* represents the total number of pixels considered.

To quantify the errors on force reconstructions, we measured the deviation from magnitude (DTM), and the deviation from angle (DTA), comparing the simulated force field and the reconstructed force field.

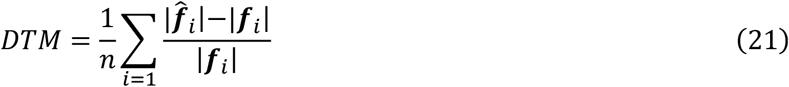

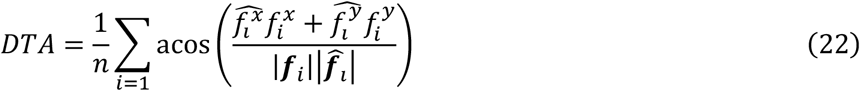

Where 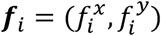 and 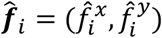 represent the simulated and reconstructed tractions for the i^th^ pixel and *n* represents the total number of pixels considered.

## 1.7 Selection of the algorithm parameters

All three techniques estimate bead displacements using a pyramidal approach—starting at low resolution and progressively refining the displacement field at full resolution. The parameters that most significantly influence the results are: the interrogation window sizes at each pyramid level for cross-correlation, the block size used for evaluating and regularizing Optical Flow, and the accumulated smoothing applied in the Demons algorithm.

To optimize these parameters, we used cell-like traction fields and evaluated performance using root mean square error (RMSE) on bead displacements, as well as the displacement-to-traction mismatch (DTM) and displacement-to-traction agreement (DTA) metrics on the traction field. Simulations were conducted for cells exerting 200, 400, and 600 Pa, with 20 replicates per condition. We also varied the point spread function (PSF) size and bead density. The results are presented in Figures S3–S11.

Among the pyramid levels, the second level had the greatest impact on cross-correlation errors. The optimal combination of interrogation window sizes—32, 16, and 8 pixels—yielded the best overall performance across all error metrics.

For the Demons algorithm, an accumulated smoothing of 2 pixels provided the best balance between accuracy and stability. Notably, the optimal smoothing increased with both bead density and PSF size.

For Optical Flow, a block size of 11 pixels offered the most favorable compromise between resolution and regularization

## 1.8 Traction force experiments

For this study, we detached the cells at the end of the timelapses and then captured the reference frame. Cells were allowed to adhere for 12 hours on TFM hydrogels mounted on 35mm glass-bottom imaging dishes (IBIDI, #**81218-200)**, followed by a 4-hour time-lapse acquisitions at the microscope. We used a temperature- and gas-controlled stage-top incubator (IBIDI, #**10720)** for imaging. We added 2 mL of 2% triton-X-100 (Millipore sigma, #**9036-19-5)** in ddH2O at 37 degrees Celsius to the 1 mL of culture medium to detach cells and capture the reference frame. We considered a Poisson ratio of 0.45 for all our experiments.

## 1.9 Gel fabrication

We fabricated polyacrylamide hydrogels following established protocols (29, 42, 43). We added a 4 µL drop of a solution containing 40% acrylamide (Bio-Rad, #161-0140) and 2% bis-acrylamide (Bio-Rad, #161-0142) to glass-bottom 35 mm dishes (IBIDI, #81218-200). Dishes had previously been activated with (3-Aminopropyl) trimethoxysilane (APTMS, Sigma-Aldrich, #281778) and a 25% glutaraldehyde solution (Electron Microscopy Sciences, #16200). To manipulate the desired rigidity, polyacrylamide mixing ratios can be found in previously published resources (29). We placed a 12 mm cover glass coated in poly-l-lysine (Sigma #25988-63-0) and dark red fluorescent micro beads (ThermoFisher, #F8807) on the drop before gels polymerized. We used tweezers to remove the coverslip, creating a TFM gel with fluorescent beads located on its surface. To prepare for cell seeding, we activated the gels using sulfo-SANPAH (ThermoFisher, #22589) and coated them with fibronectin (Millipore Sigma #FC010).

## 1.10 Cell culture

We used three cell lines in this study: MDA-MB-231 (ATCC, HTB-26), Lewis lung carcinoma (ATCC, CRL-1642) and U2OS (ATCC, HTB-96). All cell lines were cultured in Dulbecco’s modification eagle’s medium (Wicent, #319-005 CL) with added 10% Fetal bovine serum (Wisent, #090-150-FBS), 5% Penicillin-Streptomycin-Neomycin (Fisher scientific, #15640055), 5% Glutamax (Fischer scientific, #35050061).

## 1.11 Imaging

We used a Nikon Eclipse Ti2 equipped with a 20X 0.75 NA objective and a 1.5X magnification lens. The resulting pixel size on the image is 0.37 microns. Images of cells were obtained in bright field mode and those of beads with epifluorescence. We acquired images every 90 seconds for 70-80 stage positions in parallel.

## 1.12 Image analysis

We used Cellpose-SAM to segment cells from the brightfield images (44). The model generalized properly on our images with the default parameters. We then used the centroids of their masks to follow individual cells using a nearest neighbor algorithm (45). Tracks were filtered according to a minimum trajectory length, average speed, and d_max_, to avoid considering dead cells or debris.

Before computing the displacement field, all bead images were registered to the reference frame using rigid registration algorithms. The registration transformations obtained from beads images were applied to cell images as well.

Before measuring the bead displacement, we applied a Laplacian of a Gaussian, which enhances the contrast at bead locations smooths the noise.

To calculate the traction based on traction for a single cell at a specific timepoint, we only considered the force vectors located under the cell mask.

## 1.13 Definition of mechanical features

We analyse the cell using a multivariate mechanotype encompassing 6 mechanical features: trajectory persistence, speed, force coefficient of variation (force CV), effective strain, mean traction, average area.

### Trajectory persistence

The persistence of the cellular trajectory is calculated with the average of a time dependant measurement of the trajectory asymmetry. We calculated the persistence of the cellular trajectory by comparing the eigenvalues of the gyration tensor on the trajectory (46, 47).

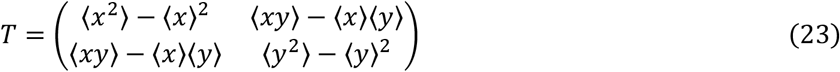

The brackets stand for a moving average. The eigenvalues of *T* are:

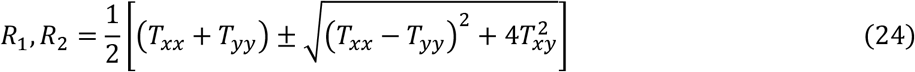

And the time dependent trajectory asymmetry parameter, that characterizes a random walk:

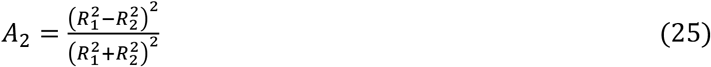

*A*_*2*_ ranges from 0 for circularly symmetric trajectories to 1 for linear trajectories.

Finally, the trajectory persistence is the time average of the trajectory asymmetry:

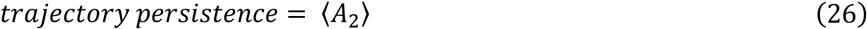

### Speed

We measure the speed as the average of instantaneous speed

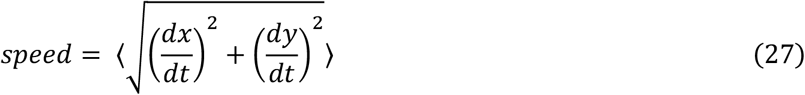

### Effective strain

After we measure and calculate the cellular tractions as described in the section 1.5, we integrate the magnitude of the force over a the cell mask for every time point. Effective strain is defined as

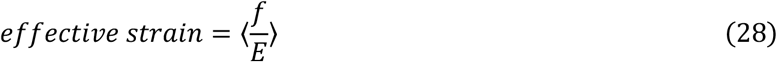

Where *f* is the total force as a function of time, *E* is the Young modulus of the substrate.

### Mean traction

The mean traction is the total force divided by the cell area and averaged through time :

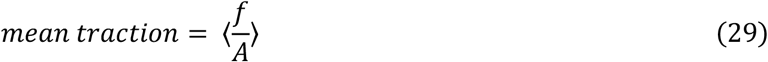

Where *A* is the area of a cell.

### Force CV

We calculate the standard deviation divided by the mean:

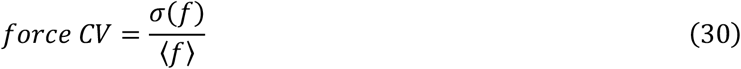

## 1.14 Statistical analysis

### 1.14.1 Effect size

We calculated the effect size as follows:

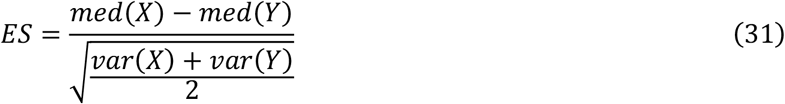

where *X* and *Y* are the two compared observations, *var* is the variance and *med* the median. We can interpret the values 0.2, 0.5, 0.8 and 1.3 as small, medium, large and very large effect sizes respectively.

## 1.15 Results

### 1.15.1 Homogeneous displacement field

We first tested the case of homogeneous displacement fields and observed that the Demons algorithm consistently outperformed all other methods in terms of displacement accuracy for a wide range of PSF sizes and bead densities. More specifically, both the error of displacement measurement (RMSE) and its dispersion are almost negligible for small displacement (Figure 1, Figure S12). Interestingly, for unresolved beads that mimic out-of-focus images with a PSF size of 10 pixels or smaller, the Demons algorithm is the only approach that allows measuring displacements over 20 pixels. This suggests that larger PSF can help to track greater displacements, and that the Demons is particularly well suited for measuring beads displacements from unresolved images. For all the techniques, an undulating pattern can be observed in the error graphs for sub-pixel bead displacements: valleys occur at full-pixel offsets, leading to a boost in correlation, and error increases for fractional displacements.

**Figure 1.**
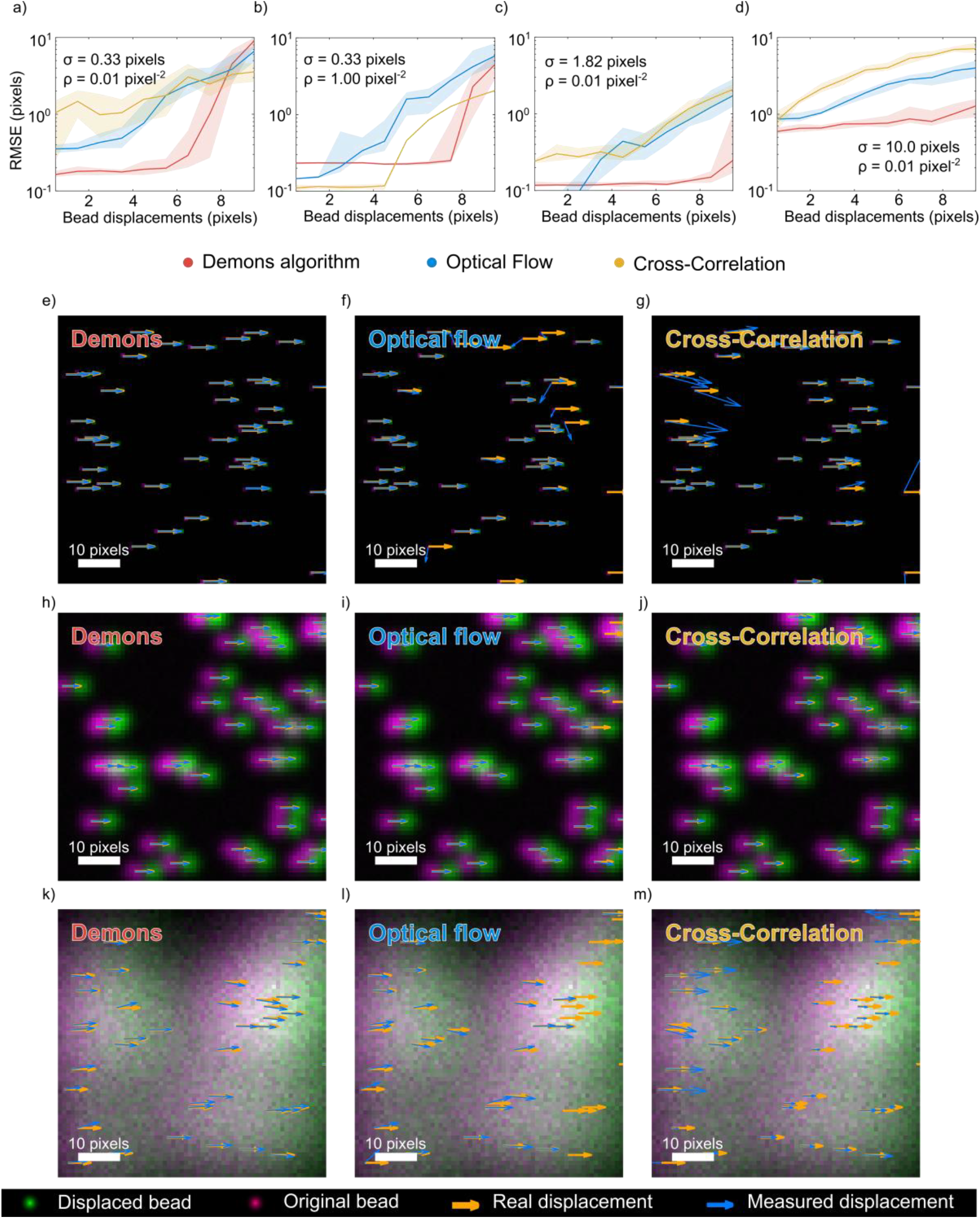
Bead displacement measurement error in homogeneous displacement fields. (a–d) Root Mean Square Error (RMSE) of bead displacement measurements. Each data point corresponds to the RMSE from a single image, with 20 replicates per displacement condition. Line thickness of the shaded regions represents the standard error of the mean. Four representative conditions are shown: (a) σ = 0.33 pixels, ρ = 0.01 beads pixel-2 (b) σ = 0.33 pixels, ρ = 1 beads pixel-2 (c) σ = 1.8 pixels, ρ = 0.01 beads pixel-2 (d) σ = 10 pixels, ρ = 0.01 beads pixel-2 (e–m) Representative bead images illustrating displacement errors for the three methods—Demons, Optical Flow, and Cross-Correlation—under the condition ρ = 0.1 beads/pixel^2^ and a displacement of 4.5 pixels (chosen for visual clarity). (e–g) σ = 0.33 pixels (h–j) σ = 1.8 pixels (k–m) σ = 10 pixels Here, σ denotes the standard deviation of the Gaussian Point Spread Function (PSF), and ρ represents bead density.

Increasing the number of beads does not significantly affect the precision of Demons estimates for small movements. The small error increases abruptly after a threshold that scales with the size of the PSF according to a complex relationship (Figure 1). This likely stem from the algorithm’s built-in smoothing mechanism, which minimizes fluctuations between adjacent vector components. Moreover, despite having no effect on the error in magnitude, raising bead densities improves angle accuracy. The mistakes of Demons are negligible for a large movement span, irrespective of the PSF’s dimensions or the number of beads, it provides consistent outcomes and remains resilient under varying imaging circumstances such as gel expansion, and nonhomogeneous bead density.

Optical Flow is often described as reliable in the case of small displacements (48); in our hands it excels at detecting sub-pixel shifts with mid-sized PFS of 1.82 pixels. However, the error and variability increase significantly when measuring displacements larger than 1 pixel. Regardless of PSF size, the error distribution of Optical Flow is worse with the increase of bead concentration, as shown in Figure S12, but a large fraction of the relative RMSE can be attributed to angular deviations, rather than the magnitude, as illustrated in Figure S3-S4. Using a well sampled 10-pixel PSF, Optical Flow struggles to maintain precise measurements, even at low bead densities, leading to an average relative RMSE between 0.2 and 0.4.

While cross-correlation tends to lag behind the other two methods, it performs the best for smaller PSF. In fact, its relative RMSE increases with PSF width, while it plateaus for increasing bead density. Across all displacement ranges, cross-correlation underestimates the values, as the magnitude error remains below 0 (Figure S3).

## 1.16 Reconstruction of cell-like force field

To evaluate the impact of displacement field errors in the force reconstruction of cellular traction, we simulated a cell pulling on a substrate. We simulated forces originating from a distribution of focal adhesions found in migrating cells (Figure 2a-c). Then, we calculated the displacements of single beads embedded in the gel using the Boussinesq approximation.

**Figure 2.**
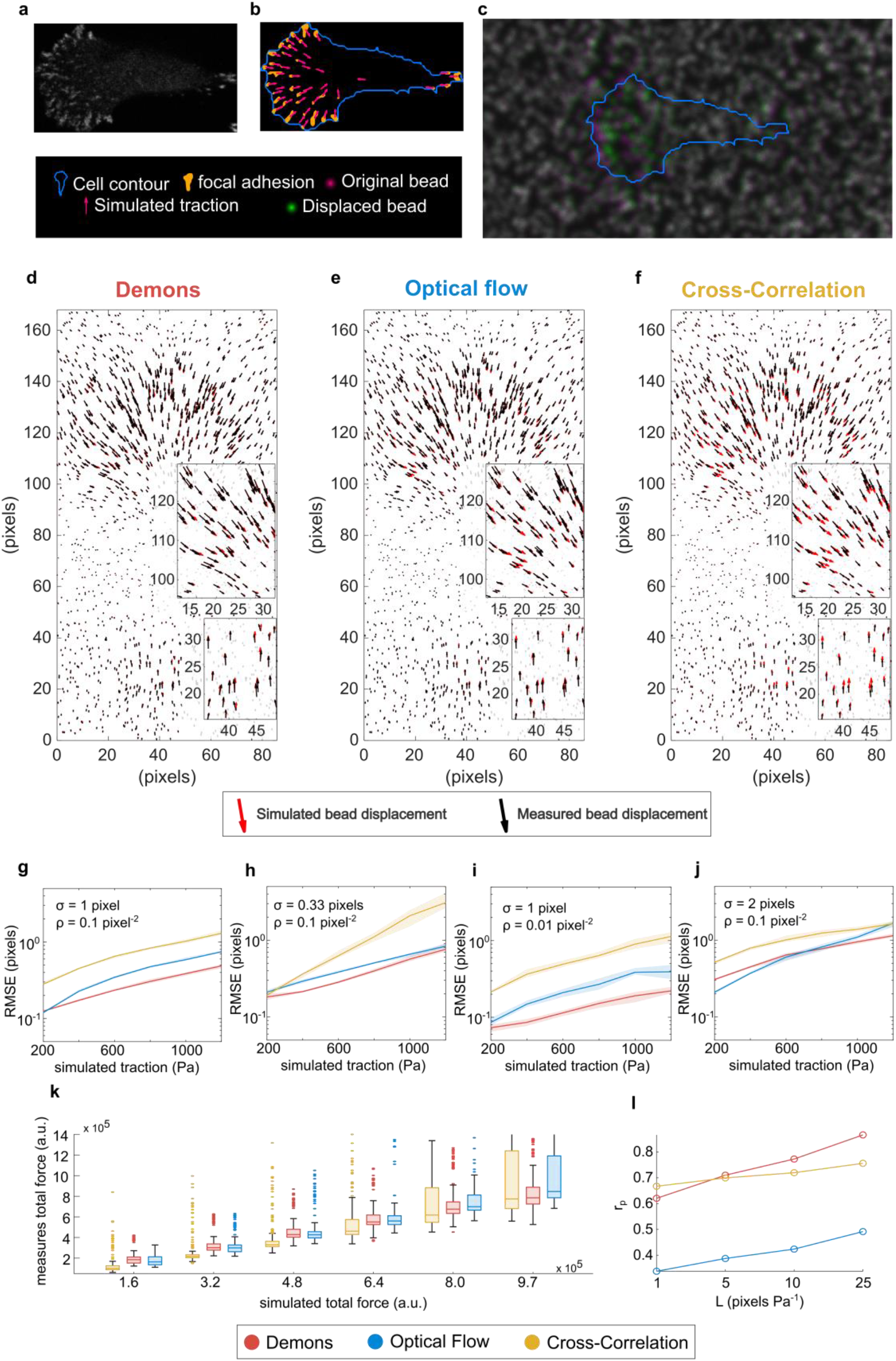
Traction force reconstruction error for a realistic cell-like bead displacement pattern. a) Confocal image highlighting focal adhesions of a migrating breast cancer cell (MDA-MB-231 expressing paxillin-mCherry). b) Segmentation of cellular membrane and focal adhesions and simulated of traction field. Cell body contour (blue), focal adhesion segmentation (orange) and simulated traction (magenta). A homogeneous traction is applied on the whole FA surface, oriented toward the centroid of the cell body. c) Simulated bead images before (magenta) and after (green) traction was applied. (d-f) Representative comparison between simulated and measured bead displacements. Every vector is associated with a single bead. For this representation, we used σ = 1 pixel, ρ = 0.1 pixel^−2^ and a traction of 600 Pa on every pixel. (g-j) Bead displacement measurement errors (RMSE) as a function of simulated traction. Thickness of curves represent percentiles 25 and 75, and the line represents the median of 200 images. 4 conditions are depicted: (g) 1-pixel PSF STD and bead density of 0.1 pixel^−2^, (h) PSF STD of 0.33 pixel and density 0.1 pixel^−2^, (i) 1 pixel PSF STD and density 0.01 pixel^−2^, and (j) PSF STD 2 pixels and density 0.1 pixel^−2^. (k-l) Prediction power of the three displacement measurement techniques. All simulated conditions are pooled together to mimic a real-life scenario. (k) Calculated total force as a function of the simulated total force. For this plot, we used a regularization parameter L=25 pixels Pa^−1^ (l) Pearson’s correlation between simulated and reconstructed total force as a function of the regularization parameter.

First, we compared the errors of displacement fields produced by Demons, cross-correlation and Optical Flow as bead density varies across the surface of a TFM hydrogel. All techniques exhibited worsening performance for increasing densities (Figure 2g-j, Figure S13). Demons displayed less variability and cross-correlation performed the worst across all bead densities. Interestingly, all techniques showed improved performance when paired with the mid-sized PSF with a standard deviation of 1 pixel. Errors nearly doubled when dealing with PSFs featuring 2-pixel standard deviations, emphasizing the crucial impact of PSF size on accuracy.

We assessed the precision of the reconstructed force field by measuring the deviation to traction magnitude (DTM) and angle (DTA). We evaluated the error for various regularization parameters (L) ranging from 1 to 25 pixels/pascal. Enhancing the regularization parameter led to an increase in DTM error, as it underestimated the force’s magnitude (Figure S14). An optimal regularization parameter is difficult to find, as there is no clear change in the slope of the L curve (data not shown). Interestingly, DTA showed a significant reduction when the regularization parameter increased, reaching over 20° difference between L=1 and L=25 (Figure S14). The relative RMSE errors reflect this trend, with the smallest error occurring at a mid-sized PSF of 1 pixel’s standard deviation. Moreover, the Demons algorithm exhibited remarkable consistency, maintaining stable DTA values for densities between 0.01 and 0.1 beads per pixel.

The characterization of heterogeneity within large sets requires not only high throughput, but also appropriate ranking of the cells. The bead tracking technique must be robust to various PSF sizes and bead densities since it is difficult to cast a perfectly flat gel and homogeneously spread beads throughout the entirety of the gel. To investigate the force ranking ability of the three techniques, we calculated the Spearman’s rank correlation coefficients between the simulated ground truth and the reconstructed force fields of simulated cells. To mimic a realistic high-throughput experiment, where the tractions of thousands of individual cells are probed over a large surface of a TFM gel, we pooled together the results for all bead densities and PSF sizes before performing the correlations. Interestingly, while increasing the regularization parameter led to lower absolute force measurements, it improved the ability to rank cells. The Demons algorithm shows a great robustness to various bead densities and PSF sizes, as it led to the best correlation and the smallest force dispersion (Figure 2l, Figure S15, Figure S16), especially for large values of traction.

We evaluated the performance of the Demons algorithm across a range of smoothing values using three key metrics: RMSE for bead displacement, deviation to traction magnitude (DTM), and deviation to traction angle (DTA). As shown in Figures S3–5, increasing the smoothing parameter consistently reduces angular error (DTA), improving directional accuracy of the reconstructed force field. However, this comes at the cost of increased DTM, indicating an underestimation of force magnitude. RMSE trends show that intermediate smoothing values yield the lowest displacement errors.

Importantly, the optimal smoothing value depends on imaging conditions: higher bead densities and larger PSF sizes shift the optimal value toward stronger smoothing. In realistic scenarios where bead density and PSF size vary across the field of view, a compromise smoothing value must be selected. Based on our results, intermediate smoothing values offer a robust balance across heterogeneous conditions, minimizing displacement error while maintaining acceptable force magnitude and angle accuracy.

### 1.16.1 High-content TFM on cell populations

To illustrate the use of the Demons algorithm in TFM on live cells, we acquired 4-hour TFM movies, for two epithelial cancer cell lines. Derived from breast cancer, MDA-MB-231 are known for their aggressive metastatic and highly migratory phenotypes, the and the U2OS are derived from osteosarcoma and are mostly still in our hands. We analyzed the mechanical properties by TFM of the two populations by seeding them on polyacrylamide gels of three distinct Young’s modulus: 6.3, 11.9 and 46.7 kilopascals.

During the TFM experiment, cell segmentation was achieved using CellposeSAM model (49), followed by automatic tracking using a Nearest Neighbor approach (45). Bead displacement fields were computed using the Demons algorithm, and we reconstructed force fields using Fourier transform traction Cytometry (39). We extracted the temporal evolution of several metrics: mean traction, affective strain, force coefficient of variation (FCV), cell area (CA), mean velocity (MV), and trajectory randomness (Figure 3 a-e).

**Figure 3.**
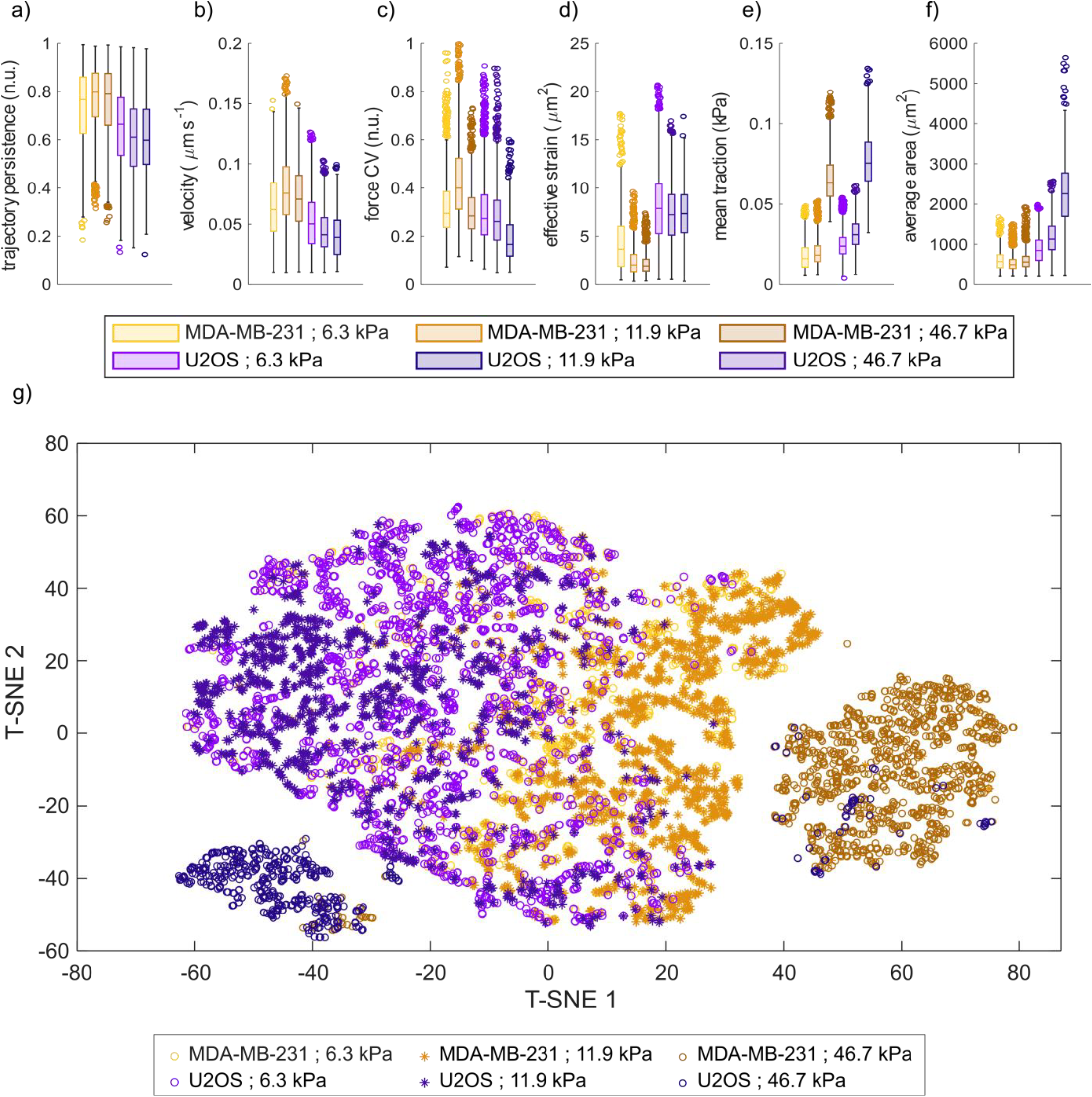
(a-f) Mechanical phenotype description using 6 features extracted from cell trajectory analysis, TFM and morphology analysis. Boxplots display distributions of single cells with median and 2 quartiles. (g) t-SNE projection of the 6 cellular mechanical features. n.u. stands for normalized units.

We categorized cells based on a multivariate mechanotype consisting of six variables. Paired variables (Figure S18) show relationships that lead to unclear segregation of cell line mechanotypes. However, dimensional reduction projections of 6-dimensional data, such as t-SNE, display differences between cell lines on different substrates (Figure 3g), which highlights the different behaviours of cells on different substrate stiffnesses. As a graphical projection of multi-dimensional data cannot fully evaluate the clusterability of datasets, we used the Dips Test, a statistical analysis that evaluates unimodality, applied to the pairwise Euclidean distances between cells using the normalized mechanical properties as coordinates (50, 51). The test refuted the assumption of unimodality for all the methods, thus we can cluster cell groups.

Furthermore, we did not observe a consistent relationship between migration speed and averaged measurement of cellular tractions, excepted effective strain, where we found a moderate Spearman correlation of 0.34 and 0.33 for MDA-MB-231 and U2OS respectively, only on the softer substrate (Table S1). However, we measured a consistent moderate correlation between migration speed and force CV, which suggests that traction dynamics is linked to migration speed.

## 1.17 Discussion

Experimental conditions and the specific scientific question determine a trade-off between computational time, complexity, cost, and accuracy to properly chose analytical tools; this work provides a thorough examination to guide such choices. Fine grained descriptions of the dynamics of single focal adhesions are often not needed to characterize large populations of cells, while large throughput data may not be informative about the details of protein interactions in migratory phenomena. The vast majority of the cell migration literature is dedicated to obtaining mechanistic molecular insight, tested on low numbers of individual cells, and only few studies aim at characterizing mechanical properties of cell types. Our work sought to analyze and compare a novel approach to obtain metrics that can be used to describe and mechanically sort populations of cells, in a cost-effective way.

Our simulations show that the bead displacement measurement accuracy on the surface of a TFM hydrogel has a significant impact on the reconstruction of cellular forces. We propose the use of Demons as an alternative to the most widely used image cross-correlation algorithms, as this method is significantly faster. We added Optical Flow to the comparative analysis, which produced great outcomes for small stresses (26). Numerical simulations demonstrated that Demons outperforms the other two alternatives and can be reliable even for displacements significantly larger than the PSF of the microscope. Demons proved to be resilient to a range of experimental conditions, particularly for displacements below 8 pixels. Optical Flow excelled only in sub-pixel displacements of beads around the pixel size, outperforming other techniques, but was the least accurate for a wide distribution of measurement errors. Cross-correlation performed optimally for PSF smaller than a pixel.

Interestingly, our findings align with observations from fluid mechanics studies, where particle tracking methods have been evaluated in similar contexts. Several studies comparing cross-correlation and Optical Flow for analyzing particle movement in turbulent flows (52–54) found that Optical Flow offers higher resolution and accuracy, achieving up to a twofold improvement over cross-correlation. While Optical Flow is more sensitive to intensity variations caused by particle motion or lighting, it excels at resolving fine displacement structures. In contrast, cross-correlation is more effective for larger displacements or lower-quality images.

Although the Demons algorithm has known limitations in low-variance regions and near image borders (55), our application to fluorescent bead images mitigates these issues. Preprocessing with a Laplacian of Gaussian filter enhances local contrast, and sufficient bead density ensures robust displacement estimation. Additionally, image regions were selected to avoid border artifacts, and the use of Efficient Second-Order Minimization (ESM) by the Demons algorithm avoids numerical instabilities associated with Hessian matrix computation (56).

We analyzed the precision of force reconstructions mimicking the traction originating from a cell. While all three techniques showed to be sensitive to changes in bead density, Demons was the most resilient. Cross-correlation performed the best with highly resolved beads, and the other two techniques worked optimally for more realistic PSFs, larger than one pixel size. Although all techniques ranked cells well according to their traction force, Optical Flow performed the worst at measuring proportional relashionship between simulated and reconstructed total force.

By combining variations in point spread function (PSF) size and bead density, the Demons algorithm consistently achieved the most accurate cell ranking based on total force. This result is particularly significant given the practical challenges of casting perfectly flat hydrogels and achieving uniform bead distribution across large surfaces—especially when working with soft gels. In high-throughput experiments, it is common to encounter out-of-focus beads and heterogeneous bead densities, making robustness to such variability a critical advantage.

Using the Demons algorithm, we successfully distinguished two cell lines across three substrate stiffness conditions by applying a two-dimensional nonlinear graphical projection of a multivariate dataset comprising six single-cell spatio-temporal mechanical properties. Furthermore, statistical analysis using the Dip Test across six cell populations rejected the unimodal distribution hypothesis, indicating the presence of underlying heterogeneity within the cancer cell population and suggesting that this approach can effectively dissect subpopulations based on mechanical behavior.

Thikonov’s method, which was used throughout this work to regularize the reconstruction of the force field, imposes a penalty on excessive norms of calculated stresses that results from inconsistencies between neighboring displacement vectors. This kind of noise is typical of cross-correlation and Optical Flow, but not Demons. Instead of using the norm of the force to regularize the reconstruction process, other operators, such as the curl or the divergence of the force field can be minimized (57). Otherwise, machine learning approaches can be employed to reconstruct the force from displacement field, as these tools are adaptable to various noise type (58, 59).

The Demons algorithm integrates regularization; it is a local iterative method that is prone to getting stuck in local minima which constrains the method to relatively small displacements (60). Additionally, its regularization process resembles a diffusion process, whereas we aim to monitor the transformations of an elastic substance. These limitations could potentially be addressed by using variations of the algorithm that are better suited for elastic materials or that better capture larger displacements (60, 61).

Current efforts are mostly aimed at enhancing 2D TFM by achieving high precision measurements of individual focal adhesions. Our primary interest, however, lies in characterizing forces across populations, thus enabling to pinpoint uncommon cells that exhibit distinct mechanical properties (reviewed in (23)). Our findings reveal that a wide-spread algorithm, that is not typically used for TFM, provides the right compromise between time and accuracy for high-content measurements of mechanical work. It resilient to experimental constrains, such as out-of-focus beads and uneven bead distribution on a large hydrogel substrates. Additionally, Demons enables the use of lower magnification objectives for increased throughput, accurately categorizes cells based on their strength, and enhances clustering capacity.

## 2 Author contributions

Nicolas Desjardins-Lecavalier designed research, performed research, analysed data, and wrote the manuscript. Santiago Costantino designed research and wrote the manuscript.

### 3 Ackowledgement

We thank to Sam Brown and Melanie Bluteau for their help in the development of experimental protocols. This work was supported by a grant from NSERC (RGPIN-2021-03330). Nicolas Desjardins-Lecavalier is recipient of BESC D scholarship from NSERC. SC holds the Wolfe Professorship in Translational Research.

