## Supporting Material for "A new approach for high-content traction force microscopy to characterize large cell ensembles"

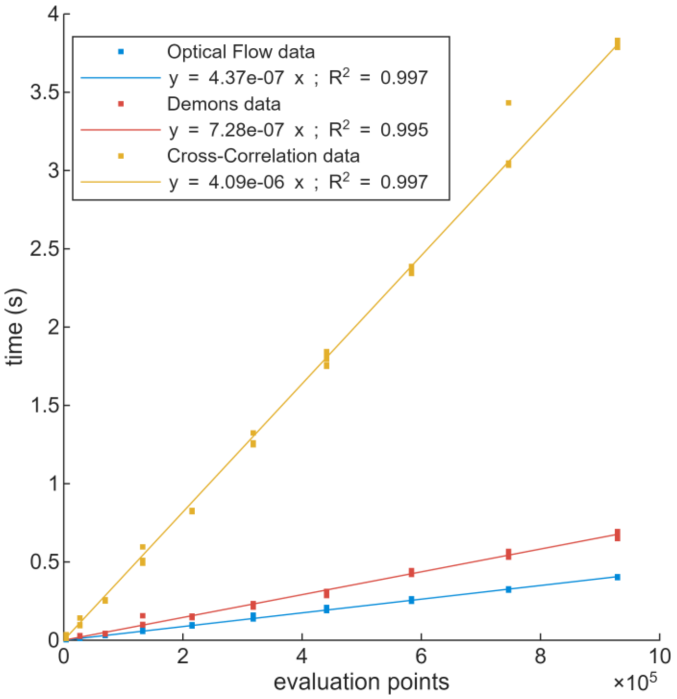


Figure S1: Computation time of the three bead displacement measurement techniques as a function of the number of evaluation points. Solid lines are spline fits on the data. Only one pyramid level is used for Cross-Correlation, with a window size of 16x16 pixels, and a linear sampling every 4 pixels. Demons used 15 iterations, while Optical Flow used 5 iterations. Computations were made on a Intel Core i9 – 10850k processor.

### Estimation of the number of floating-point operations

Reviewers #1 and #3 raised concerns regarding the efficiency comparison between the algorithms. Although we provided computation times in Supplementary Figure 1, we acknowledge that runtime can vary significantly depending on implementation details. To complement the runtime analysis, we now include an estimation of the number of floating-point operations (FLOPs). While a complete FLOP count for pyramidal algorithms is challenging, we provide estimates for a single pyramid level across all techniques. Additionally, we report the number of iterations required to stabilize the bead displacement error for both the Demons algorithm and Optical Flow. Details follow :

#### Optical Flow

The Optical Flow algorithm minimizes the brightness constancy constraint by aligning a warped version of the moving image, denoted as $I(W(x;p))$, with the template image $T(x)$.

$$\sum_{\boldsymbol{x}} \left[\boldsymbol{T}(\boldsymbol{x}) - \boldsymbol{I}\big(\boldsymbol{W}(\boldsymbol{x};\boldsymbol{p})\big)\right]^2$$

Where the warp transform is simply the translation of pixels located at *x* by *p*

$$W(x;p) = \begin{bmatrix} x_1 + p_1 \\ x_2 + p_2 \end{bmatrix}$$

This minimization is performed with respect to the displacement parameters *p*, and constitutes a non-linear optimization problem. The optimization follows a steepest descent approach, requiring the computation of the image gradient **∇*I*** and the Jacobian ***δW/δp***. A complete description of the algorithm is accessible elsewhere (15). The next table gives an estimation of the FLOPs for each steps of the algorithm, for a single iteration. The number of iterations required to converge are displayed at the end of this section.

| # | ALGORITHM STEPS | Estimated FLOPs | remarks |
|---|---|---|---|
| 1 | Wrap ***I*** using ***W(x;p)*** to compute ***I(W(x;p))*** | 12 N | Bilinear interpolation |
| 2 | Compute error image ***T(x)*** - ***I(W(x;p))*** | N | One subtraction per pixel |
| 3 | Compute the gradient ***∇I*** (***W(x;p)***) | 36 N | Convolution with 3×3 kernel (e.g., Sobel), two channels |
| 4 | Evaluate Jacobian δ***W*** / δ***p*** at ***(x;p)*** | 76 N | Includes convolutions and derivative divisions |

| | | | |
|---|---|---|---|
| 5 | Compute the steepest descent images $\nabla I \ \ \delta W / \delta p$ | 6 N | Matrix multiplication per pixel |
| 6 | Compute Hessian matrix $H = \sum_x [\nabla I \ \ \delta W / \delta p]^T [\nabla I \ \ \delta W / \delta p]$ | 3 N | Element-wise multiplication and summation |
| 7 | Compute update term $\sum_x [\nabla I \ \ \delta W / \delta p]^T [T(x) - I(W(x;p))]$ | 2 N | Two multiplications per pixel |
| 8 | Compute $\Delta p = H^{-1}$*update term | 5/3 $N^{3/2}$ + 2 N | Matrix inversion via Cholesky decomposition |
| 9 | Update parameters $p \leftarrow p + \Delta p$ | 2 N | Simple addition per pixel |

Total Estimated FLOPs per Iteration : 140 N + 5/3 $N^{3/2}$

#### Demons Algorithm

The Demons algorithm minimizes a correspondence energy with respect to the displacement field u*u*, representing pixel-wise velocity vectors ($u=dp/dt$). In this algorithm, as mentioned previously, Efficient Second-Order Minimization (ESM) is used instead of Gauss-Newton approximation (2). Briefly, since $\nabla T \approx \nabla I(s_{optimal})$ in image registration, a second-order approximation can be used without explicitly computing the Hessian matrix.(2). Hense, the updated of an updated transform is done in the direction of the symmetric Jacobian, $J^p = -(\nabla T + \nabla (I(s)))/2$, as expressed by u :

$$u(p) = -\frac{T(p) - I(s(p))}{||J^p||^2 + \frac{\sigma_i^2(p)}{\sigma_x^2}} J^{p^T}$$

Where *I* is a moving image that we want to warp to a template image *T* with the transform *s(p)*. To ensure diffeomorphic properties of the transformation, the updated mapping *s*(*p*) is composed with the exponential of the displacement field *u*. A complete description of the algorithm can be found elsewhere (16).

| # | Algorithm Step | Estimated FLOPs | Remarks |
|---|---|---|---|
| 1 | Warp $I(s(p))$ | 12 N | Bilinear interpolation |
| 2 | Calculate T(p) – I(s(p)) | N | One subtraction per pixel |
| 3 | Compute symmetric Jacobian $J^p = -(\nabla T + \nabla(I(s)))/2$ | 75N | Four convolutions with 3×3 kernels, addition and division |
| 4 | Minimize correspondence energy $E^{corr}$ $u(p) = -[(T(p)-I(s(p))) / (\|\|J^p\|\|^2 + \sigma_i^2(p) / \sigma_x^2)] J^{pT}$ | 6 N | Three multiplications, two divisions, one addition |
| 5 | Fluid regularization $u \leftarrow K_{fluid} \star u$ | 144 N | Two convolutions with 6×6 kernels |

| | | | |
|---|---|---|---|
| 6 | Compose diffeomorphic transform $c \leftarrow s \circ exp(u)$ | 32 N | One exponentiation, one bilinear interpolation |
| 7 | Diffusion-like regularization $s \leftarrow K_{diff} \star c$ | 144 N | Two convolutions with 6x6 kernels |

Total Estimates FLOPs per Iteration : 414N

#### Cross-Correlation

Cross-correlation methods rely on template matching to estimate the transformation between a stationary and a moving image. A template is extracted from the moving image and convolved with the stationary image, producing a correlation matrix. The location of the maximum value in this matrix indicates the displacement of the template. To achieve sub-pixel accuracy, a Gaussian fit is applied to the correlation matrix, allowing precise localization of the peak displacement.

Imagine a template window of dimensions $T \times T$. To create a correlation map for a single pixel, a linear sampling rate of 1/b, we need $T^2/b^2$ correlations. The number of FLOPs required for a Fast Fourier Transform (FFT) depends on the specific algorithm used. In general, FFTs have a computational complexity of $O(n\log n)$, where $n$ is the size of the input sequence. In this case, we will consider the Radix-2 Cooley-Tuckey algorithm, which requires $5\,T\,log2(T)$ FLOPs for a $T \times T$ image. To construct the cross-correlation map $cc = ifft(fft(A) * fft(B))$, 3 FFTs are needed and one matrix multiplication, resulting in $15\,T\,log2(T) + T^2$ FLOPs. For subpixel localization, a 3-point Gaussian fit is performed on the cross-correlation matrix. Hence, the subpixel shift is expressed

$$\delta_x = \frac{\ln(cc_{-1,0}) - \ln(cc_{+1,0})}{2[\ln(cc_{-1,0}) - 2\ln(cc_{0,0}) + \ln(cc_{+1,0})]}$$

$$\delta_x = \frac{\ln(cc_{0,-1}) - \ln(cc_{0,+1})}{2[\ln(cc_{0,-1}) - 2\ln(cc_{0,0}) + \ln(cc_{0,+1})]}$$

We estimate approximately 20 FLOPs per logarithmic computation, with the total cost of sub-pixel peak localization amounting to roughly 210 FLOPs.

The total amount of FLOPs can be approximated by *[ T2/b2 * (15 T log2(T) + T2 ) + 210 ] N*

For a template window of $T$=16 pixels and a linear sampling rate of $b$=4 pixels, the total estimated FLOPs amount to approximately 19,666$N$.

#### Number of iterations

As illustrated in the following figure, the Optical Flow algorithm converges in fewer iterations compared to the Demons algorithm. We repeated the RMSE calculation five times between the estimated and ground-truth displacement fields across multiple iterations, using various bead configurations under both homogeneous and cell-like displacement conditions. As shown in the figure below, the structure of the displacement field has minimal influence on the convergence behavior. With a single pyramid level, Optical Flow typically converges within 5 to 10 iterations, while with three pyramid levels, convergence is achieved in 3 to 5 iterations. In contrast, the Demons algorithm requires 100 to 200 iterations for a single pyramid level, and 10 to 15 iterations when using three pyramid levels.

*Figure X. Convergence behavior of bead displacement measurement algorithms.*

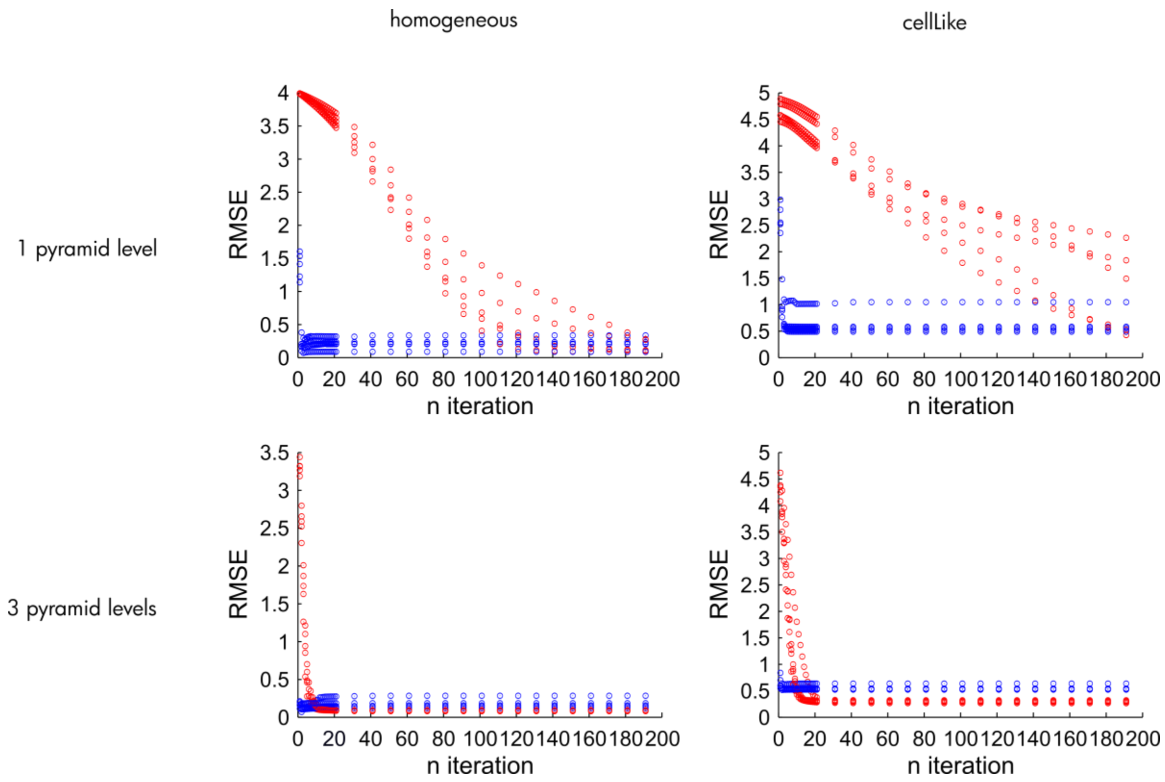


Figure S2: Convergence behavior of bead displacement measurement algorithms. The four scatter plots illustrate the evolution of RMSE values across iterations for different algorithms and pyramid configurations. Red and blue data points represent RMSE measurements under homogeneous and cell-like displacement fields, respectively. The Optical Flow algorithm demonstrates faster convergence compared to the Demons algorithm, regardless of the displacement field structure.

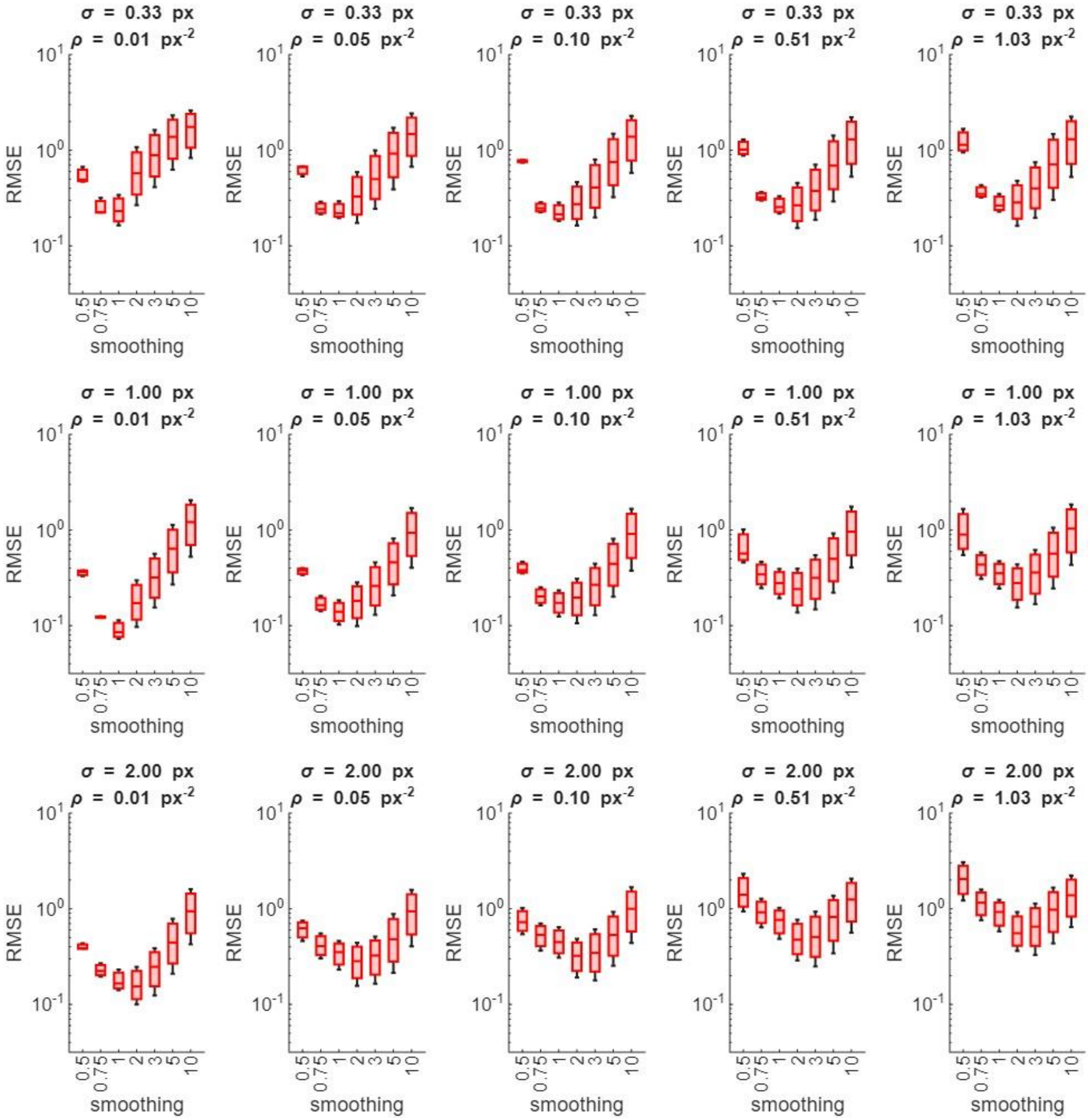

Figure S3: Demons algorithm optimization: Accumulated smoothing parameter. RMSE of simulated cell-like bead displacement pulling at 200, 400, and 600 Pa. Each traction was replicated 20 times. Boxplots show medians and quartiles. Measurements taken for 3 Gaussian point spread functions (σ) and 5 bead densities (ρ).

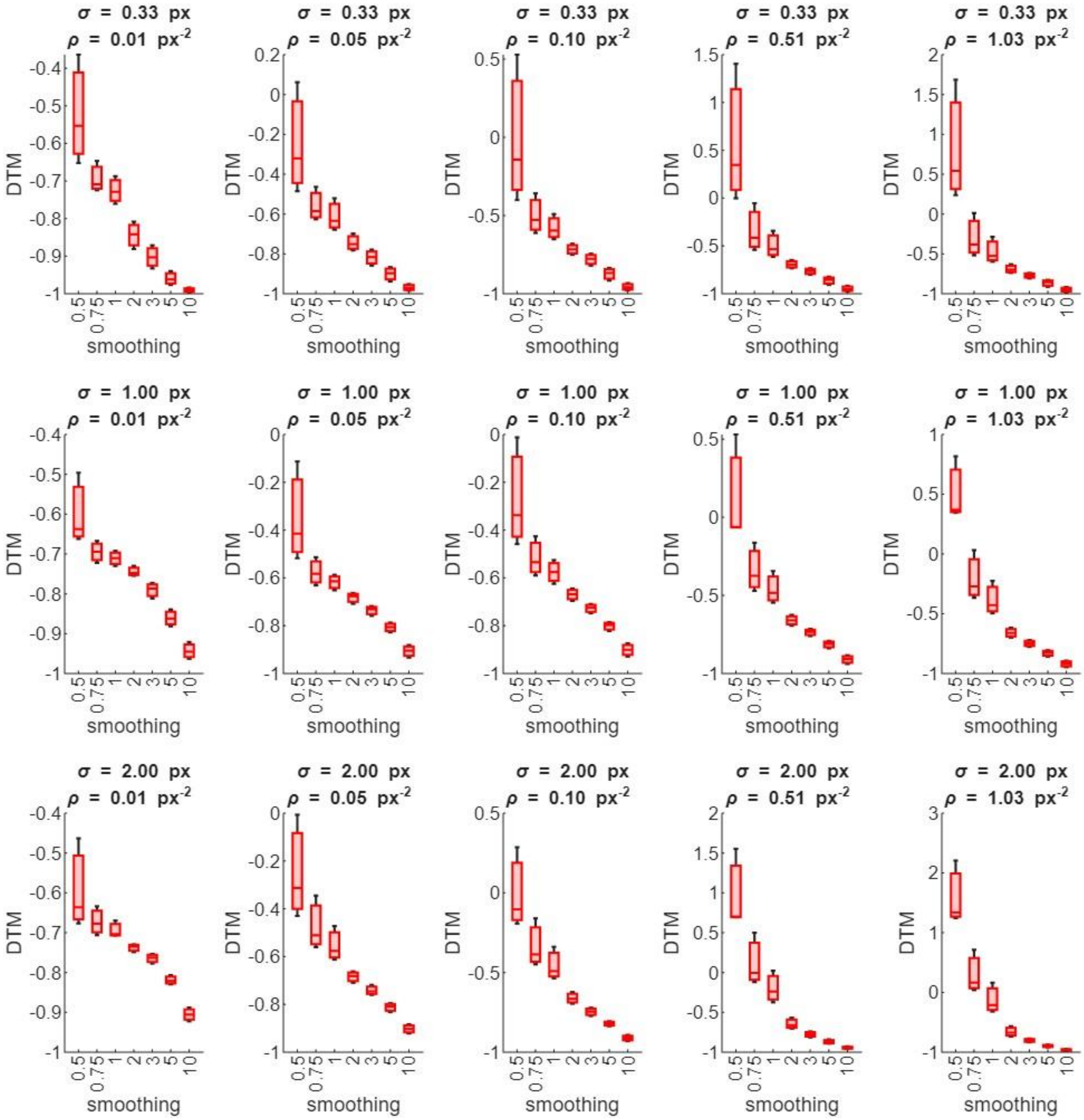


Figure S4: Demons algorithm optimization: Accumulated smoothing parameter. DTM of simulated cell-like traction forces of 200, 400, and 600 Pa. Each traction was replicated 20 times. Boxplots show medians and quartiles. Measurements taken for 3 Gaussian point spread functions (σ) and 5 bead densities (ρ).

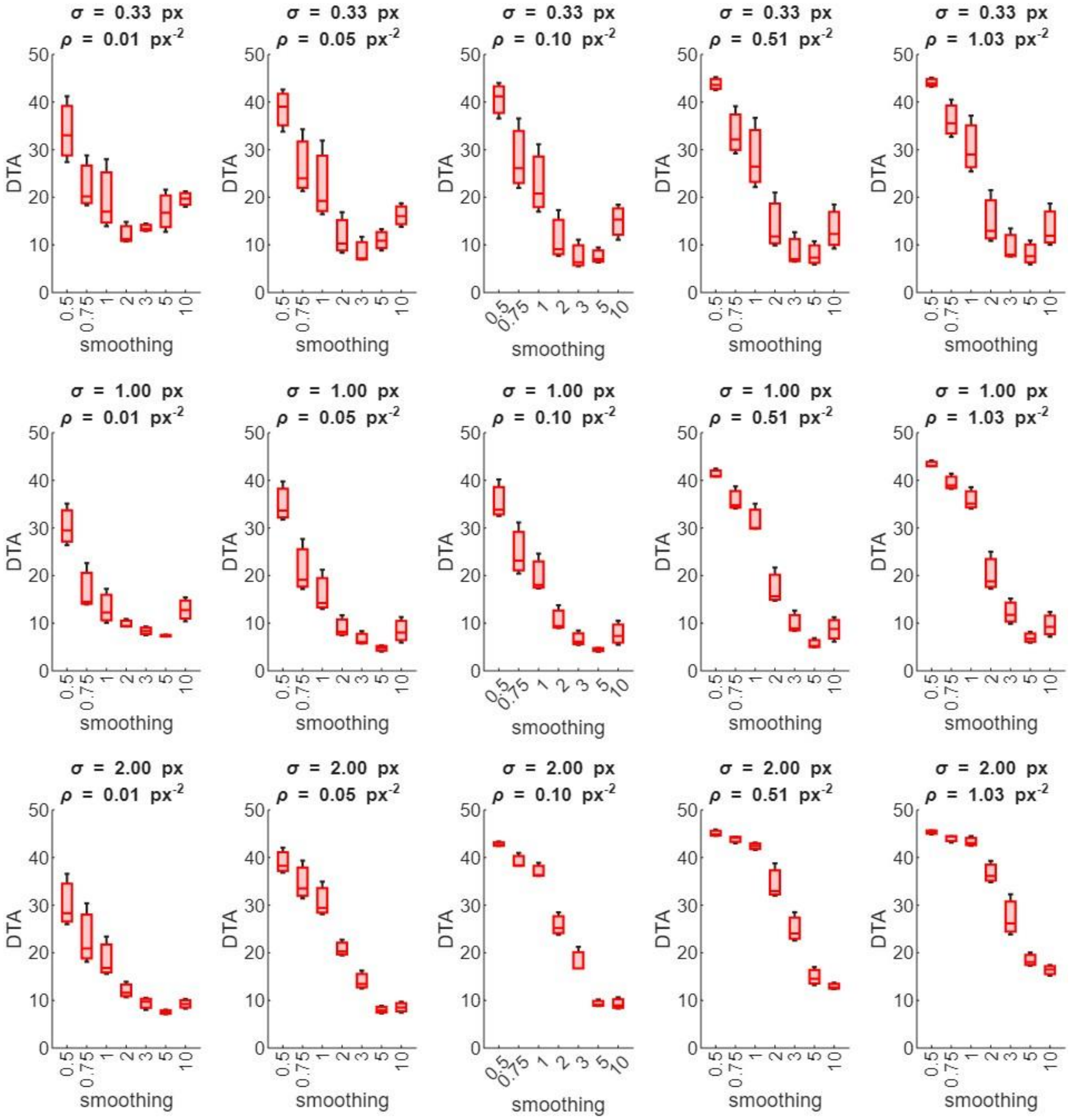


Figure S5: Demons algorithm optimization: Accumulated smoothing parameter. DTA of simulated cell-like traction forces of 200, 400, and 600 Pa. Each traction was replicated 20 times. Boxplots show medians and quartiles. Measurements taken for 3 Gaussian point spread functions (σ) and 5 bead densities (ρ).

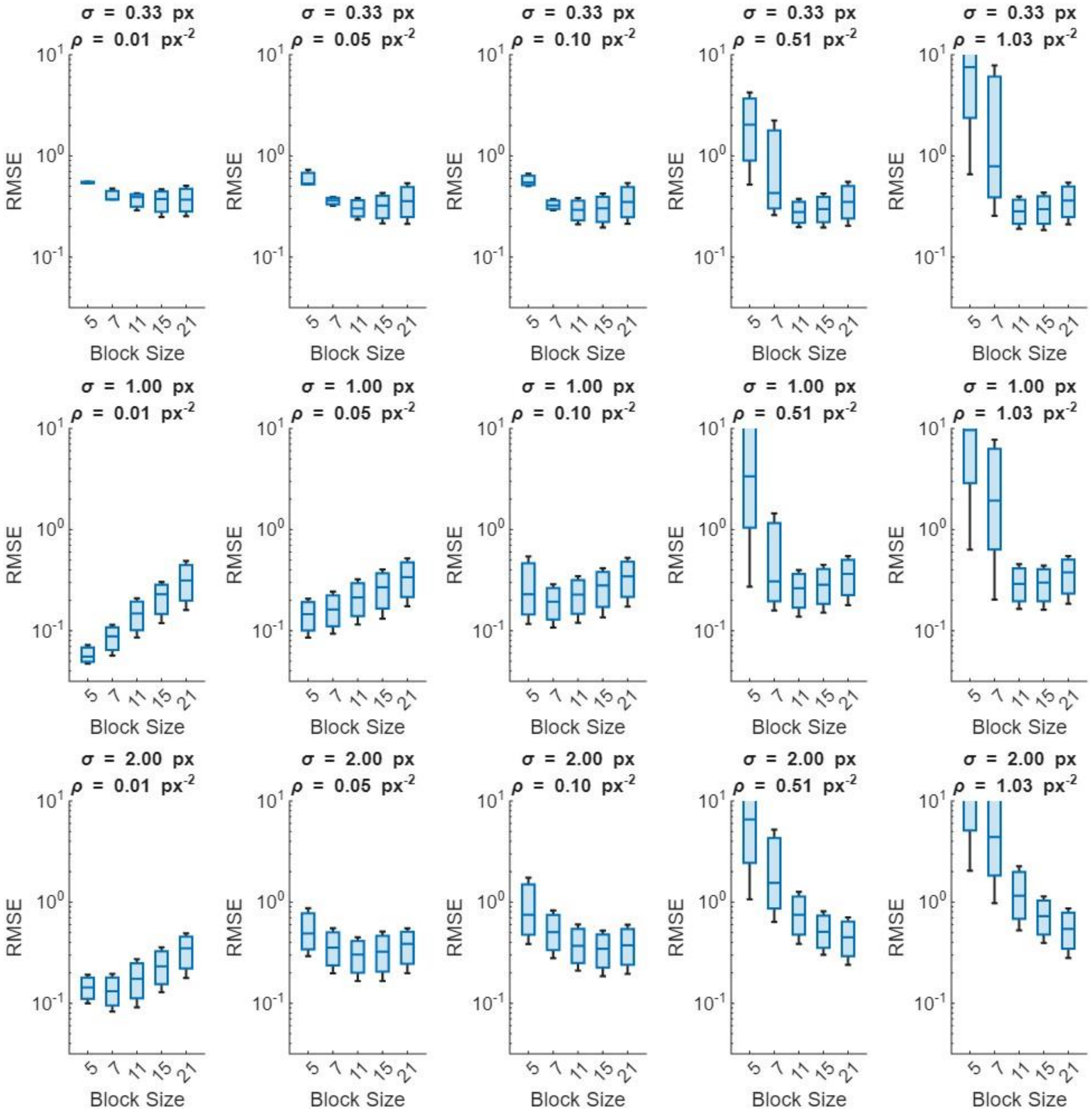


Figure S6: Optical Flow optimization: Block Size. RMSE of *simulated cell-like* bead displacement *pulling at* 200, 400, and 600 Pa. *Each traction was replicated 20 times*. Boxplots show medians and quartiles. Measurements taken for 3 Gaussian point spread functions (σ) and 5 bead densities (ρ).

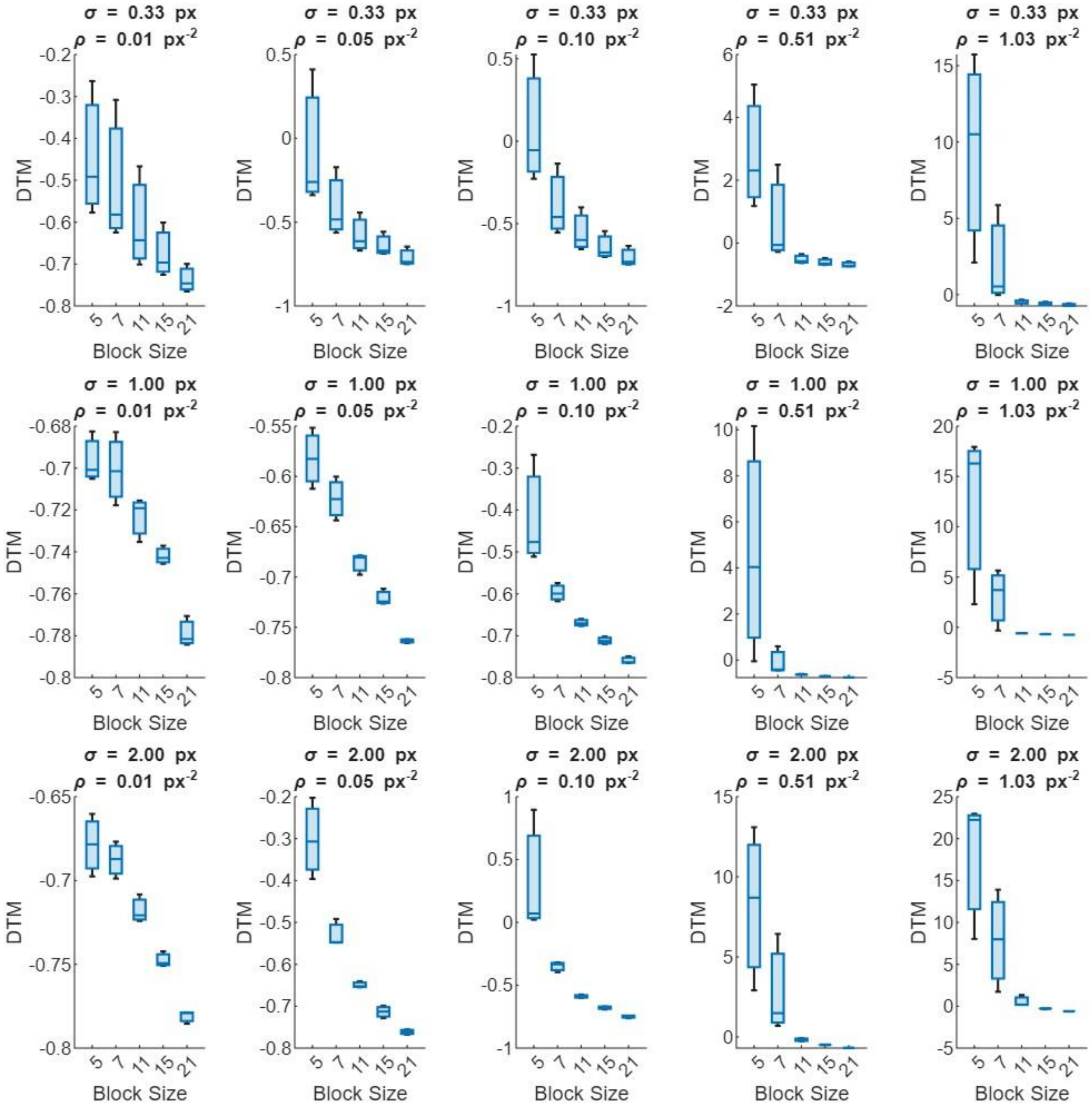


Figure S7: Optical Flow optimization: Block Size. DTM of simulated cell-like traction forces of 200, 400, and 600 Pa. Each traction was replicated 20 times. Boxplots show medians and quartiles. Measurements taken for 3 Gaussian point spread functions (σ) and 5 bead densities (ρ).

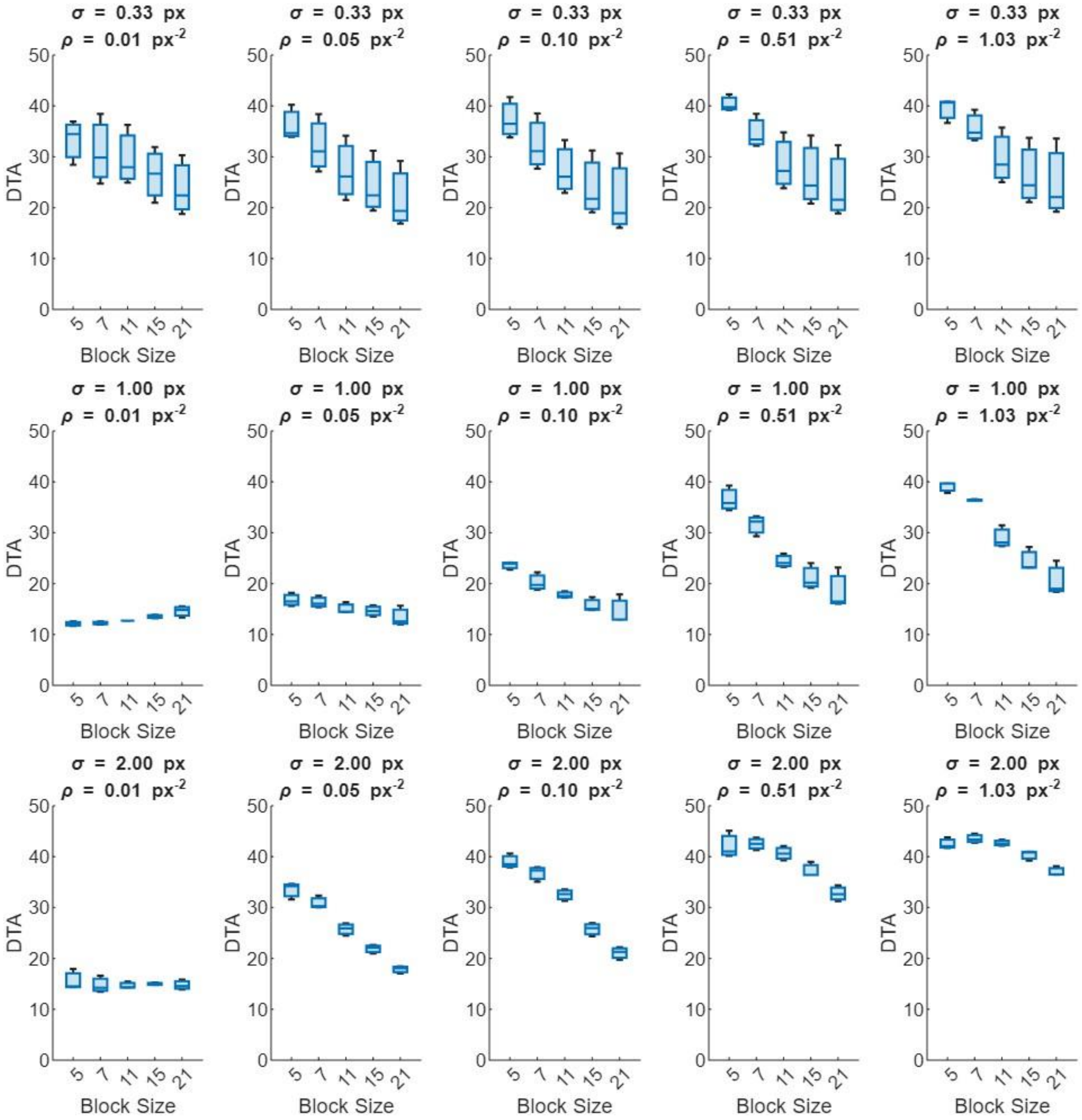

Figure S8: Optical Flow optimization: Block Size. DTA of simulated cell-like traction forces of 200, 400, and 600 Pa. Each traction was replicated 20 times. Boxplots show medians and quartiles. Measurements taken for 3 Gaussian point spread functions (σ) and 5 bead densities (ρ).

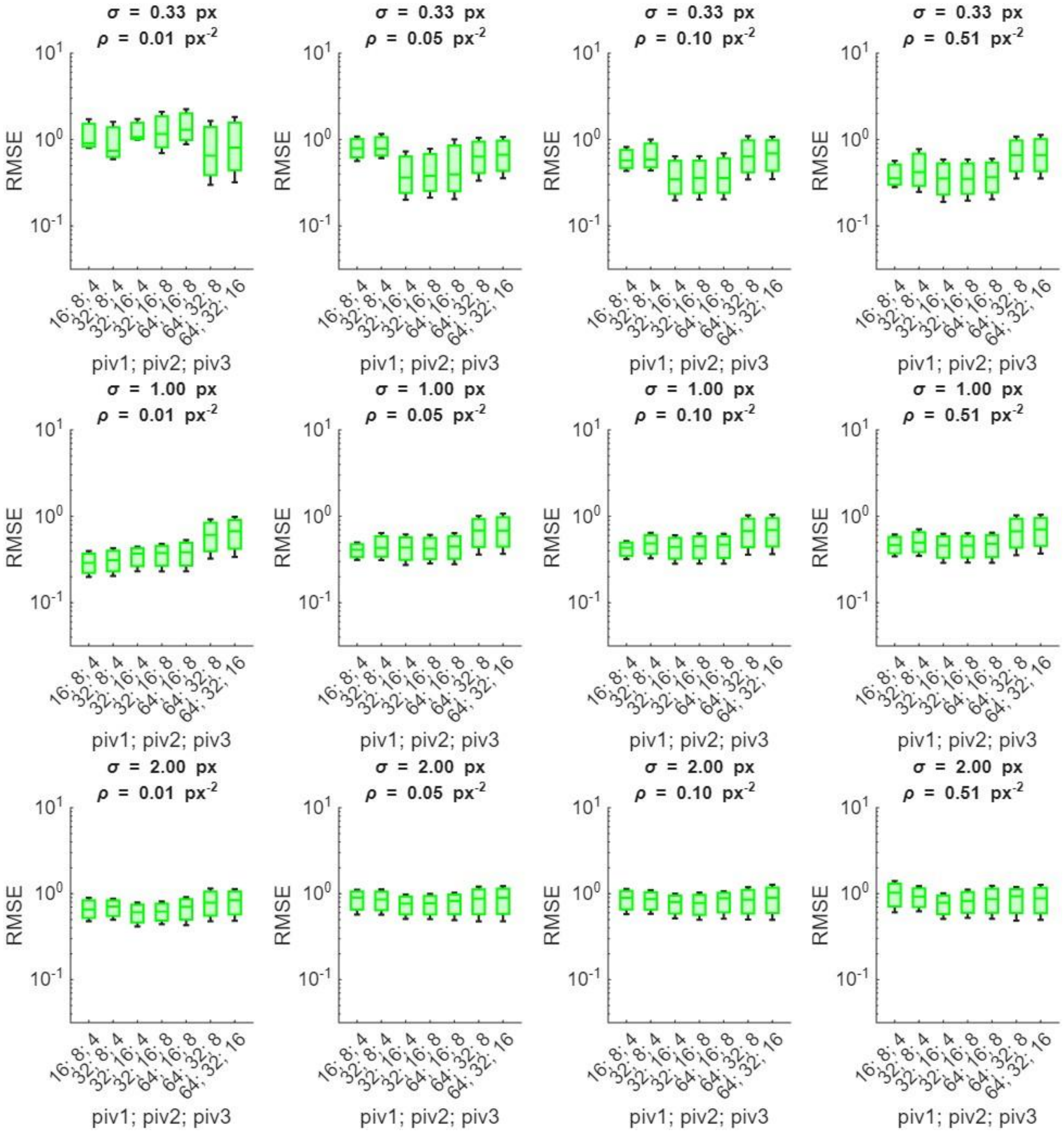


Figure S9: Cross-correlation optimization: Investigation window sizes for the 3 pyramid levels. The three window sizes are labeled *piv1; piv2; piv3*. RMSE of *simulated cell-like* bead displacement *pulling at* 200, 400, and 600 Pa. *Each traction was replicated 20 times*. Boxplots show medians and quartiles. Measurements taken for 3 Gaussian point spread functions (σ) and 5 bead densities (ρ).

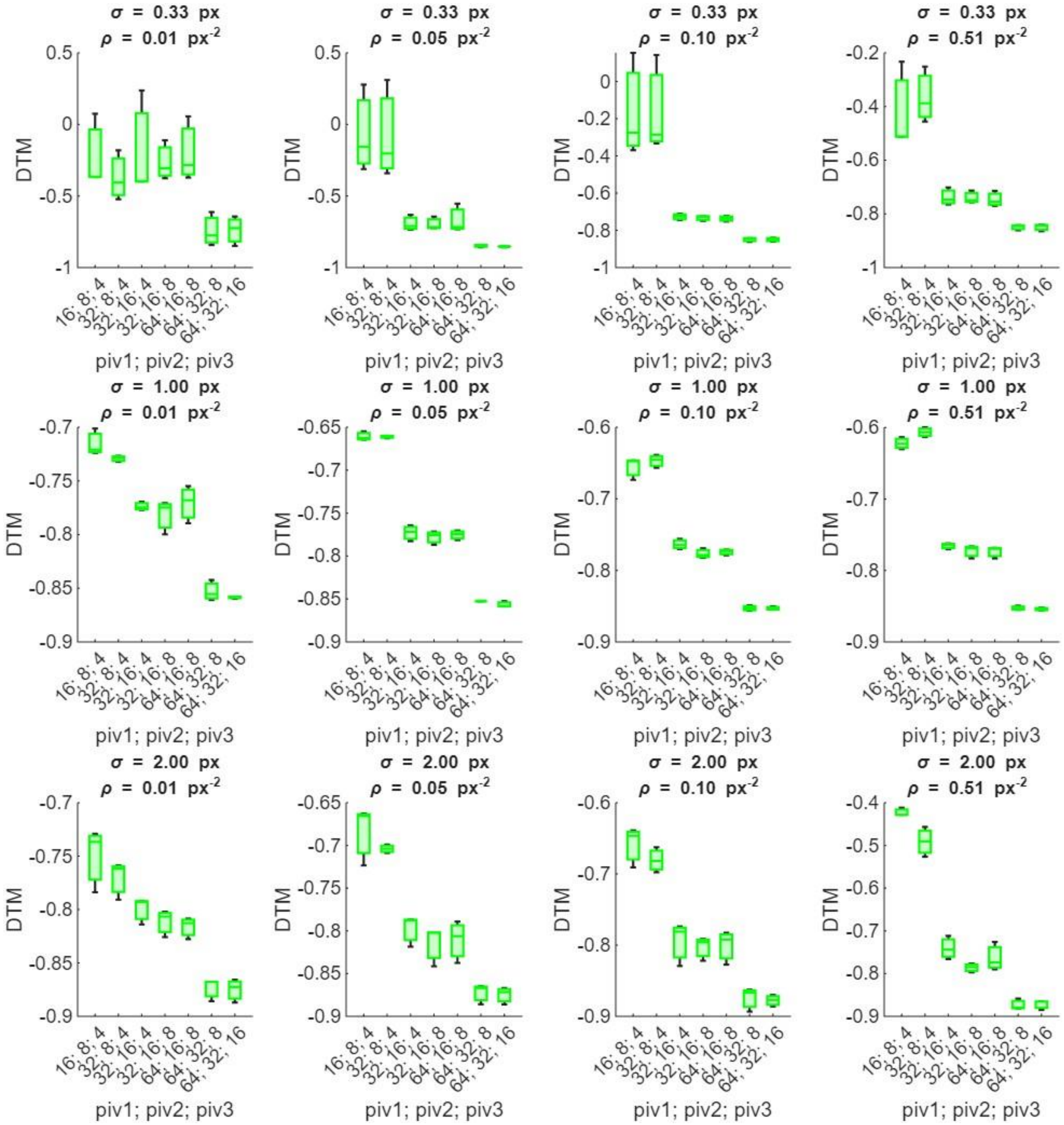

Figure S10: Cross-correlation optimization: Investigation window sizes for the 3 pyramid levels. The three window sizes are labeled *piv1; piv2; piv3*. DTM of simulated cell-like traction forces of 200, 400, and 600 Pa. Each traction was replicated 20 times. Boxplots show medians and quartiles. Measurements taken for 3 Gaussian point spread functions (σ) and 5 bead densities (ρ).

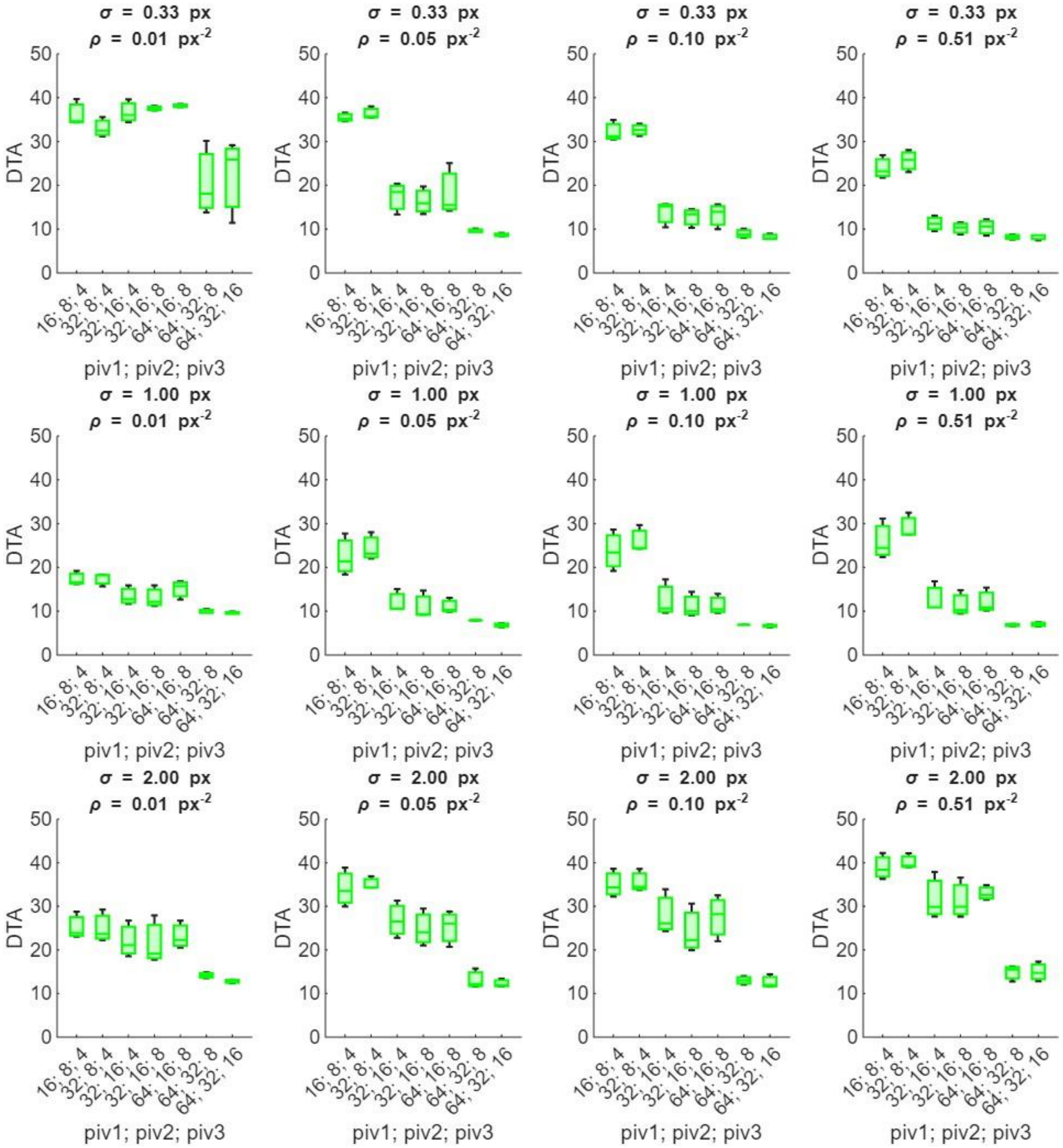


Figure S11: Cross-correlation optimization: Investigation window sizes for the 3 pyramid levels. The three window sizes are labeled *piv1; piv2; piv3*. DTA of simulated cell-like traction forces of 200, 400, and 600 Pa. Each traction was replicated 20 times. Boxplots show medians and quartiles. Measurements taken for 3 Gaussian point spread functions (σ) and 5 bead densities (ρ).

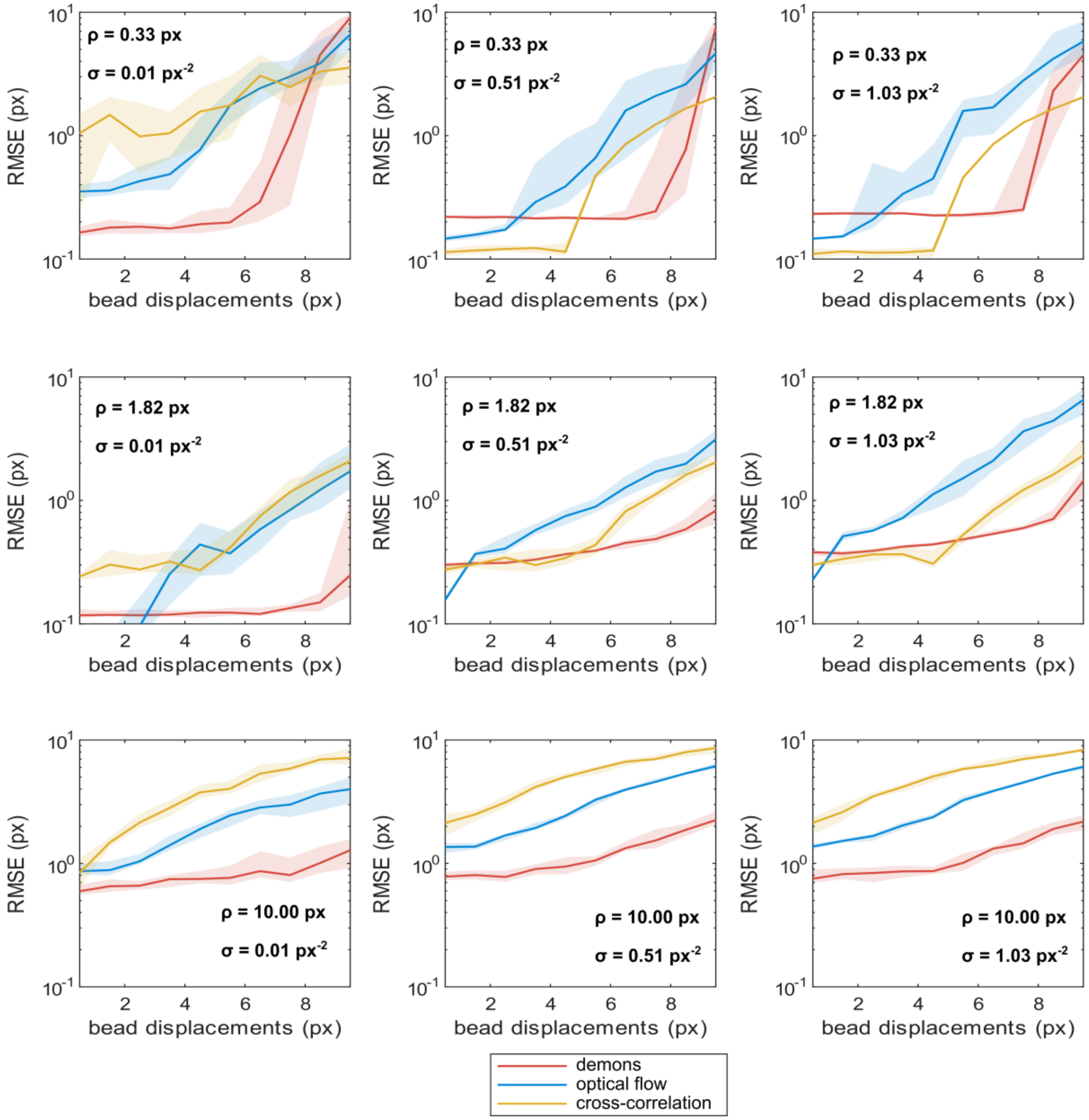


Figure S12: Displacement measurement error (RMSE) for all simulated conditions of homogeneous bead movement. σ stands for the standard deviation of the gaussian PSF, ρ is the bead density. We performed 20 images per simulated displacement magnitude, bead density and PSF size, by randomly redistributing the bead locations. Curves represent the median error and quartiles.

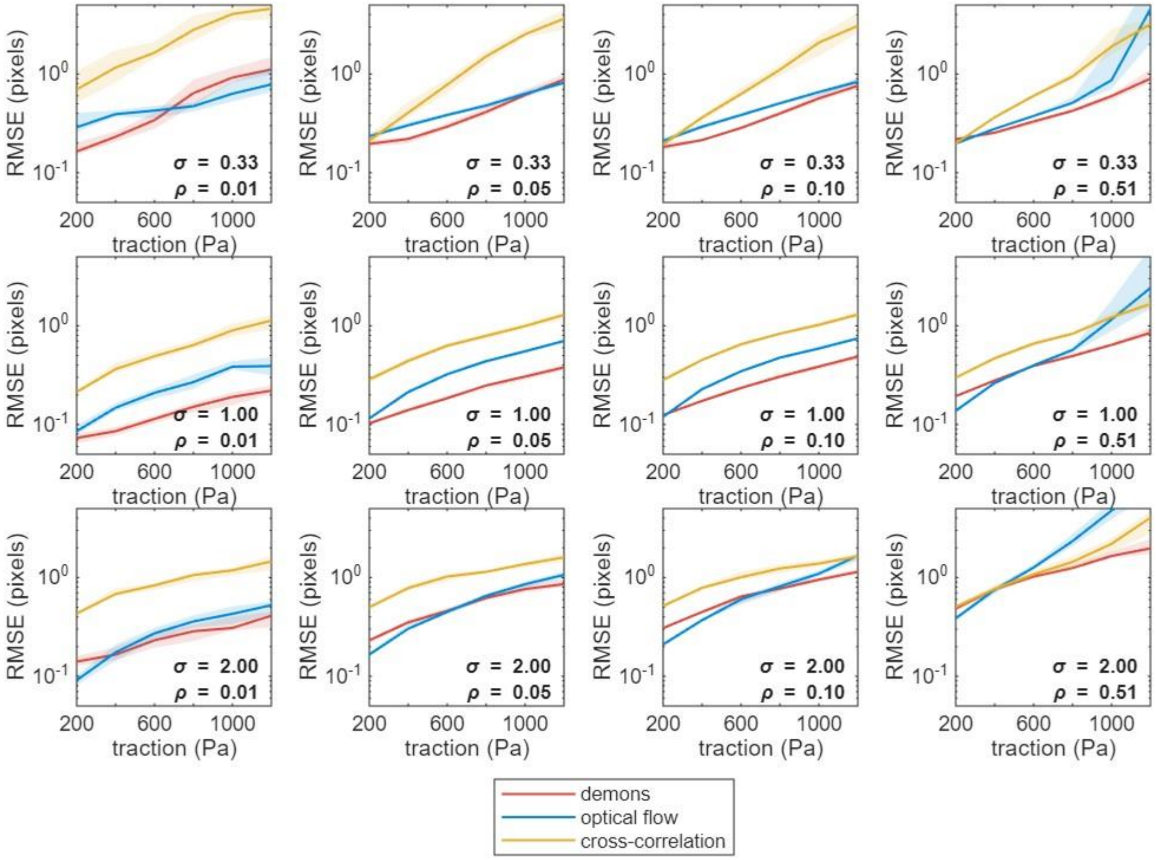


Figure S13: Displacement measurement error (RMSE) for all simulated conditions of bead movement generated by a virtual cell. σ stands for the standard deviation of the gaussian PSF, ρ is the bead density. We performed 50 images per simulated displacement, bead density and PSF size, by randomly redistributing the bead locations. Curves represent the median error and the thickness, the percentiles 25 and 75.

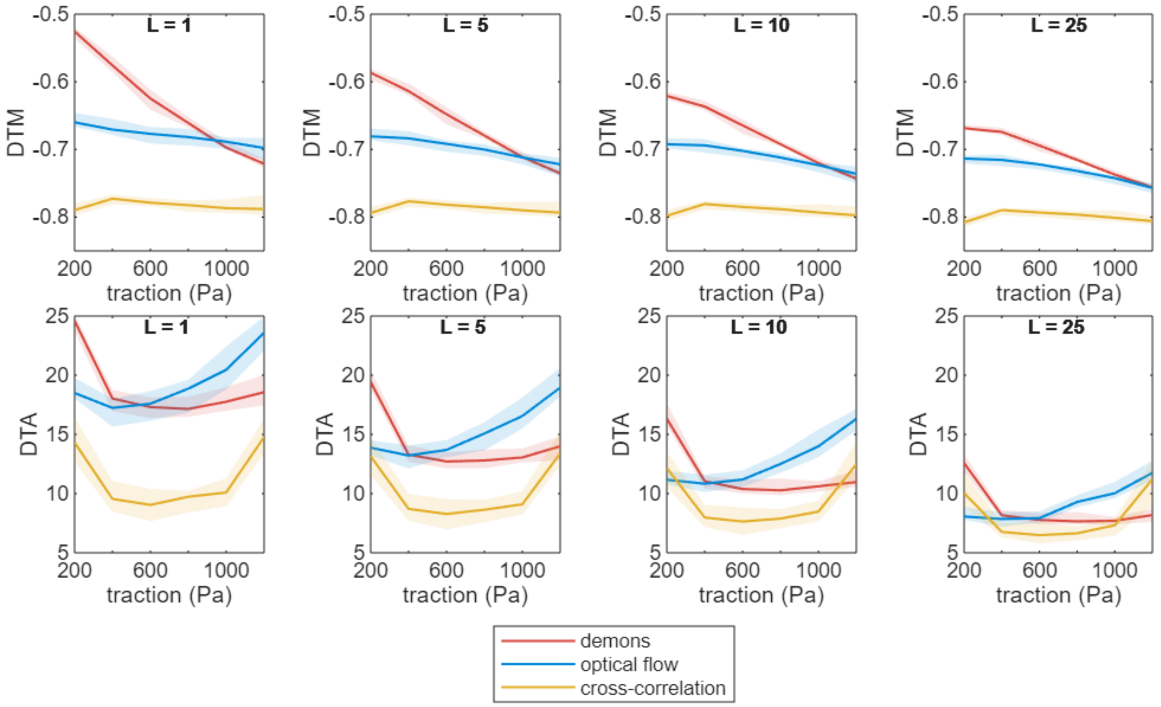


Figure S14: Impact of regularization on the quality of force reconstruction. A) Deviation to traction magnitude (DTM) and B) deviation to traction angle (DTA) of a simulated cell for bead density 0.1 pixels$^{-2}$ and PSF standard deviation 1 pixel. of a simulated cell. σ stands for the standard deviation of the gaussian PSF, ρ is the bead density. Curves in A and B represent median error and the thickness of the curve represents the percentiles 25 and 75.

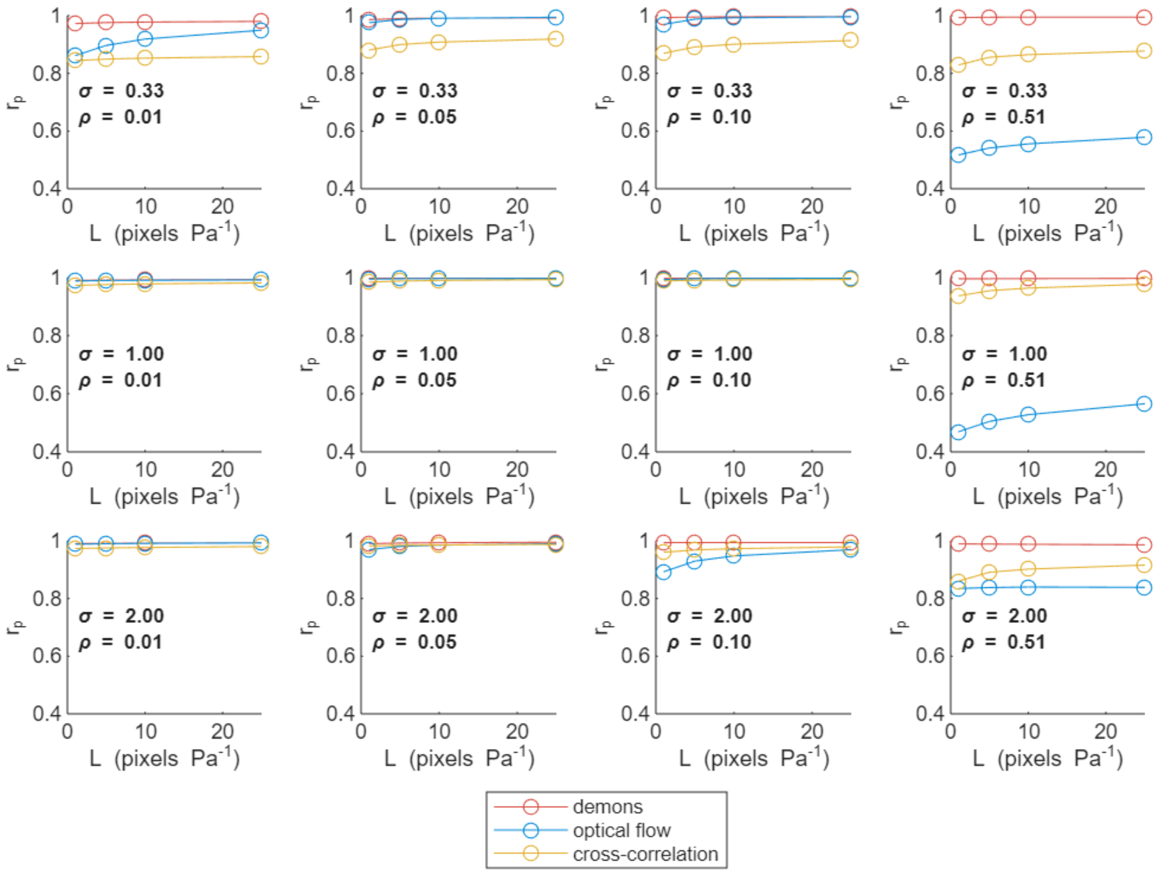


Figure S15: Ranking ability after force reconstruction: Pearson's correlation between simulated and reconstructed total force of a virtual cell as a function of the regularization parameter (L). σ stands for the standard deviation of the gaussian PSF, ρ is the bead density. Thickness of the curve represents 95% confidence interval lower and upper bounds.

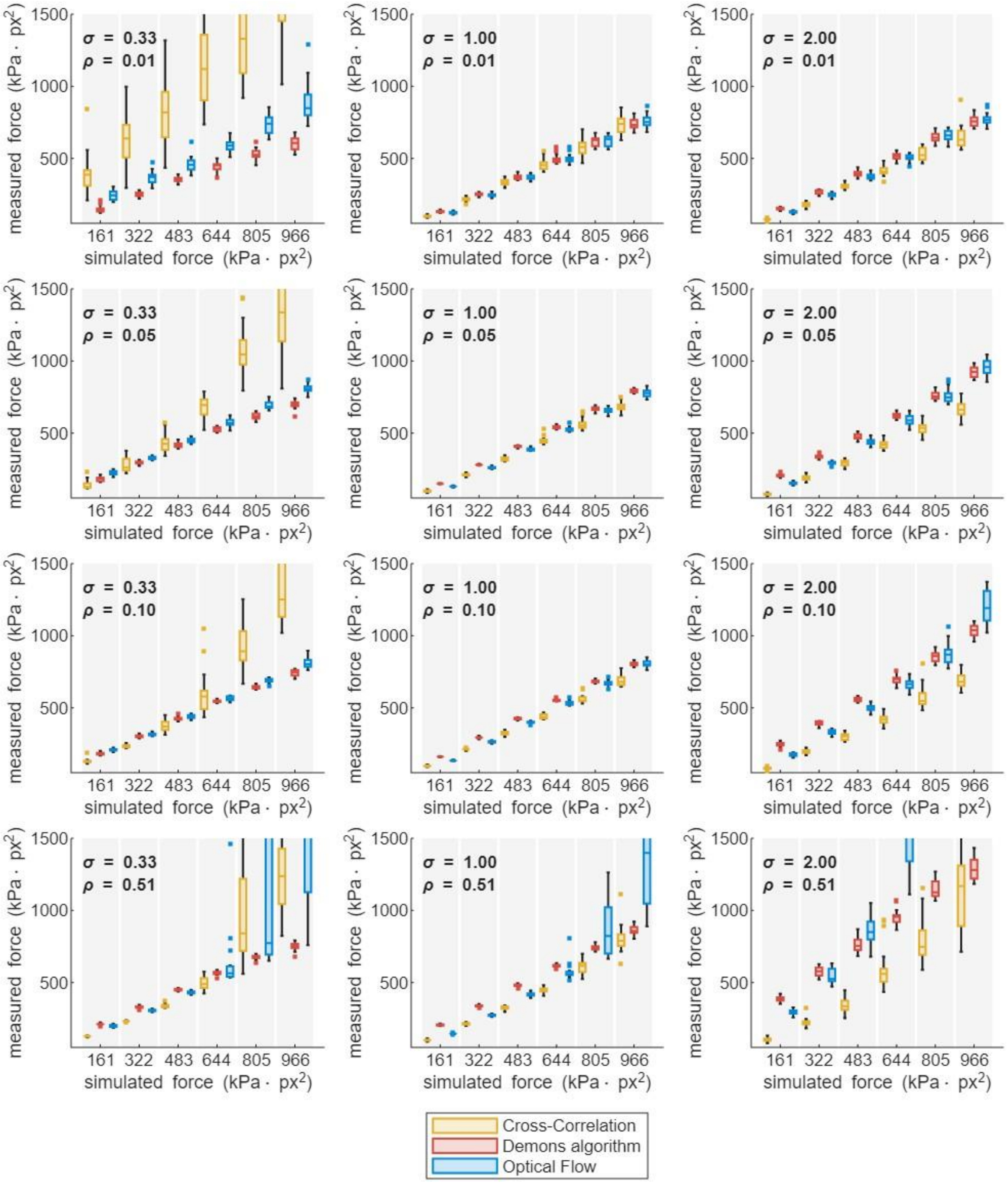


Figure S16: Prediction power of the bead tracting techniques in multiple bead density and PSF size conditions. Both simulated and measured forces are integrated over the cell mask. σ stands for the standard deviation of the gaussian PSF, ρ is the bead density. The regularization parameter is L = 25 Pa pixel$^{-1}$

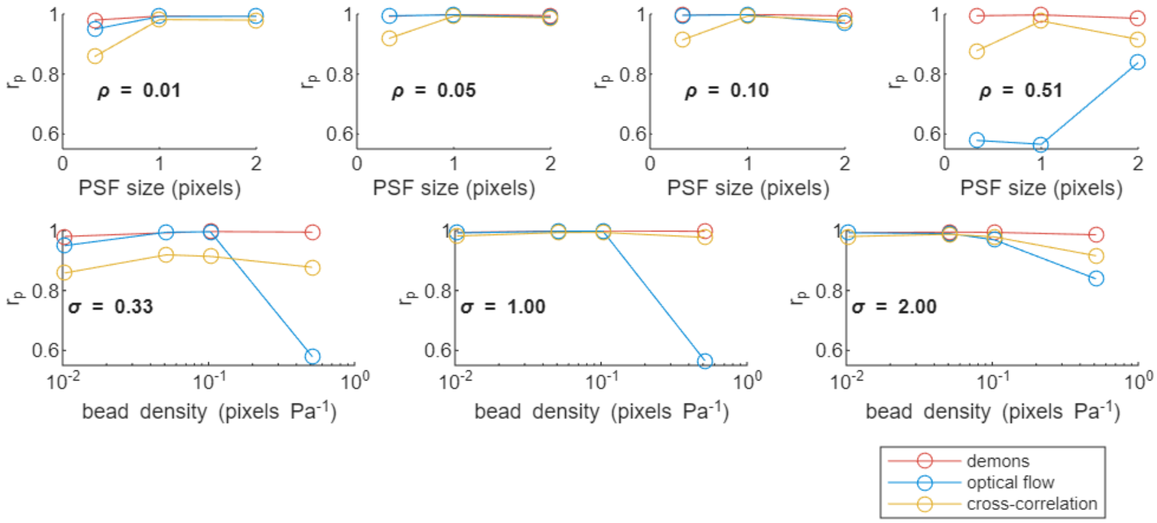


Figure S17: Ranking ability of the bead tracking techniques for multiple bead densities and PSF sizes. Pearson's correlation comparing the simulated and calculated force. σ stands for the standard deviation of the gaussian PSF, ρ is the bead density. The regularization parameter is L = 25 Pa $pixel^{-1}$

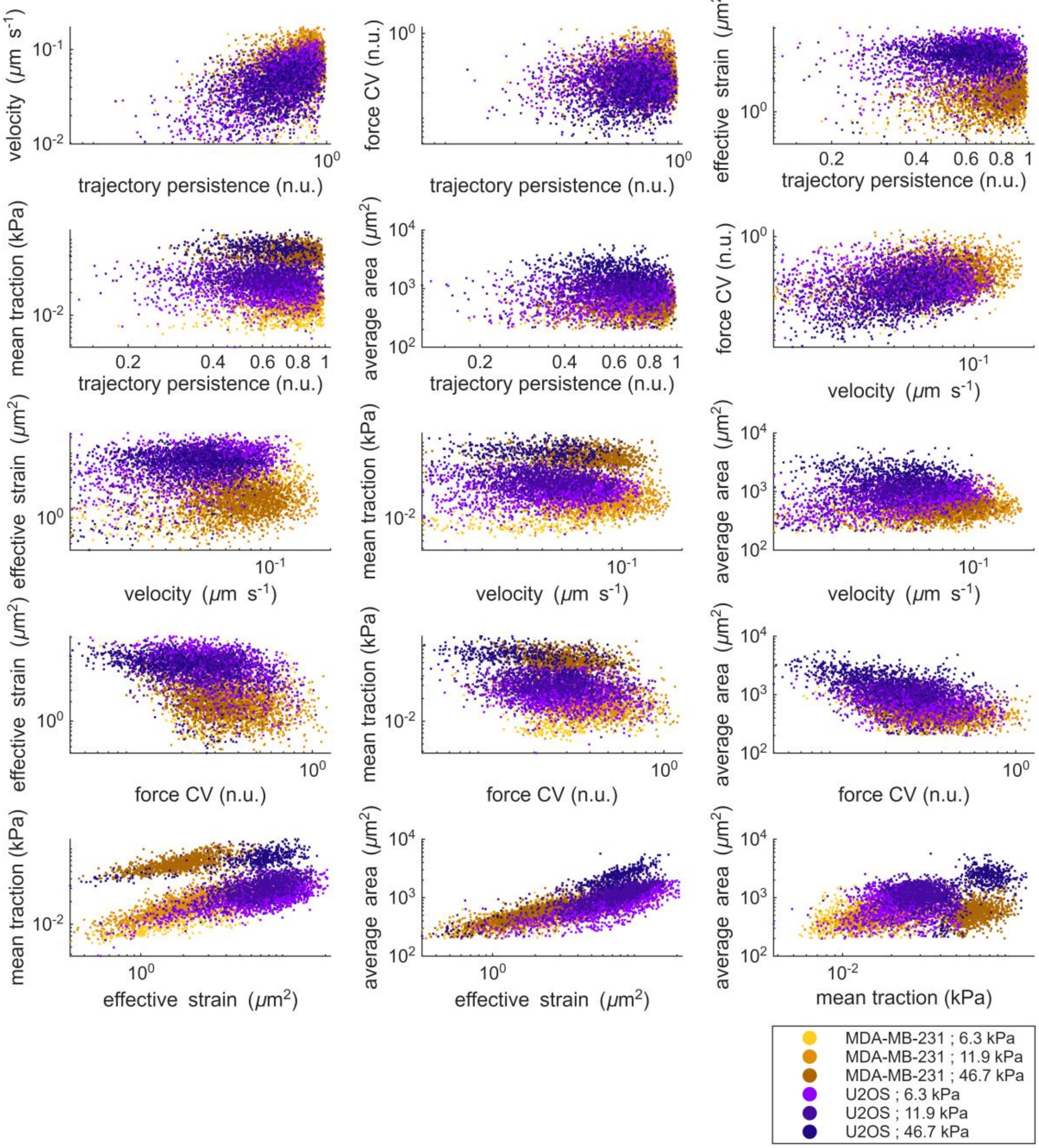


Figure S18: Distributions of 6 cellular mechanical features plotted one against another for MDA-MB-231 and U2OS on 3 polyacrylamide gel stiffnesses. n.u. stands for normalized units.

| | Trajectory randomness | Trajectory randomness | Trajectory randomness | Trajectory randomness | Trajectory randomness | velocity | velocity | velocity | velocity | force CV | force CV | force CV | effective strain | effective strain | mean traction |
|---|---|---|---|---|---|---|---|---|---|---|---|---|---|---|---|
| | velocity | force CV | effective strain | mean traction | average area | force CV | effective strain | mean traction | average area | effective strain | mean traction | average area | mean traction | average area | average area |
| MDA-MB-231 ; 6.3 kPa | 0,2 | -0,01 | 0,04 | -0,08 | 0,16 | 0,3 | 0,34 | 0,3 | 0,32 | -0,28 | -0,16 | -0,32 | 0,88 | 0,79 | 0,48 |
| MDA-MB-231 ; 11.9 kPa | 0,23 | -0,02 | 0,11 | 0,01 | 0,16 | 0,22 | 0,02 | -0,02 | 0,11 | -0,31 | -0,16 | -0,28 | 0,83 | 0,78 | 0,38 |
| MDA-MB-231 ; 46.7 kPa | 0,3 | 0,01 | 0,14 | -0,06 | 0,19 | 0,24 | 0,08 | -0,1 | 0,18 | -0,41 | -0,24 | -0,4 | 0,68 | 0,84 | 0,28 |
| U2OS; 6.3 kPa | 0,5 | -0,05 | 0,14 | -0,05 | 0,19 | 0,05 | 0,33 | 0,03 | 0,41 | -0,37 | -0,13 | -0,38 | 0,58 | 0,77 | 0,06 |
| U2OS; 11.9 kPa | 0,31 | 0,01 | -0,02 | -0,04 | 0 | 0,34 | 0,08 | -0,09 | 0,16 | -0,33 | -0,14 | -0,33 | 0,52 | 0,73 | -0,03 |
| U2OS; 46.7 kPa | 0,29 | 0,18 | -0,19 | -0,05 | -0,21 | 0,43 | -0,04 | 0,08 | -0,04 | -0,5 | -0,28 | -0,48 | 0,54 | 0,85 | 0,18 |

Table S1: Spearman ranked correlation ($r_s$) between time averaged features and p values (p) within cell lines

| | **cell line** | **stiffness** |
|---|---|---|
| **effective strain** | 1914,041 | 170,1111 |
| **mean traction** | 532,2828 | 7547,007 |
| **velocity** | 1328,545 | 31,55938 |
| **force CV** | 217,1982 | 140,7739 |
| **A2** | 5,270375 | 7,9E-208 |
| **area** | 379,7884 | 0 |

Table S2 : Anova F numbers comparing the effect of the cell line and stiffness on the 6 mechanical features.

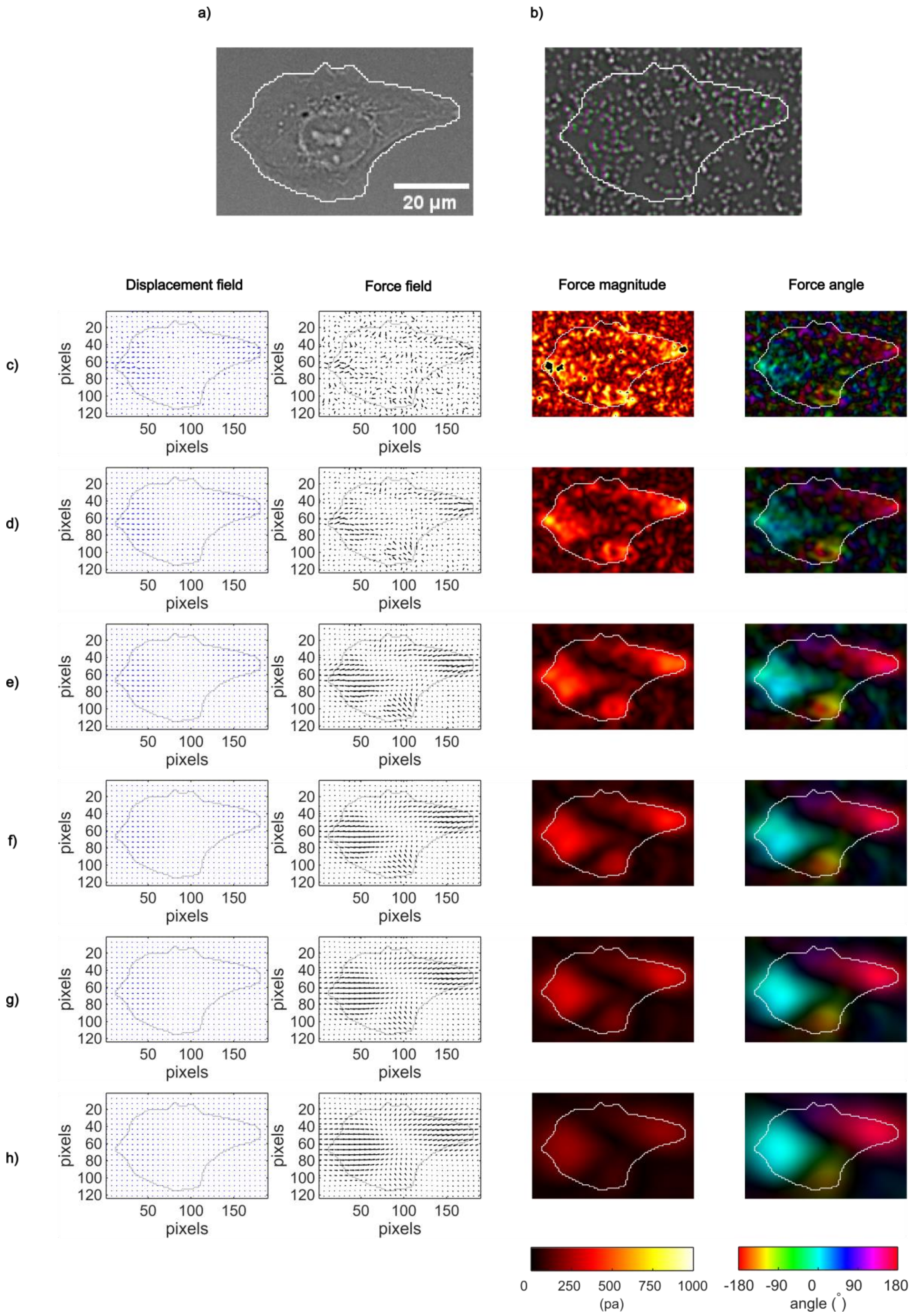


Figure S19: Effect of demons smoothing on force reconstruction for a strongly

pulling U2OS cell on a soft substrate (6.7 kPa).
(a) Brightfield image of a U2OS cell with segmentation overlay.
(b) Fluorescent bead images before (magenta) and after (green) traction, both processed with a Laplacian of Gaussian filter. Cell segmentation is shown in white.
(c–h) Displacement fields, reconstructed force fields (adjusted size of arrows for representation), force magnitudes, and force angles for various accumulated smoothing sizes: (c) 0.75, (d) 2, (e) 4, (f) 6, (g) 10, (h) 12.

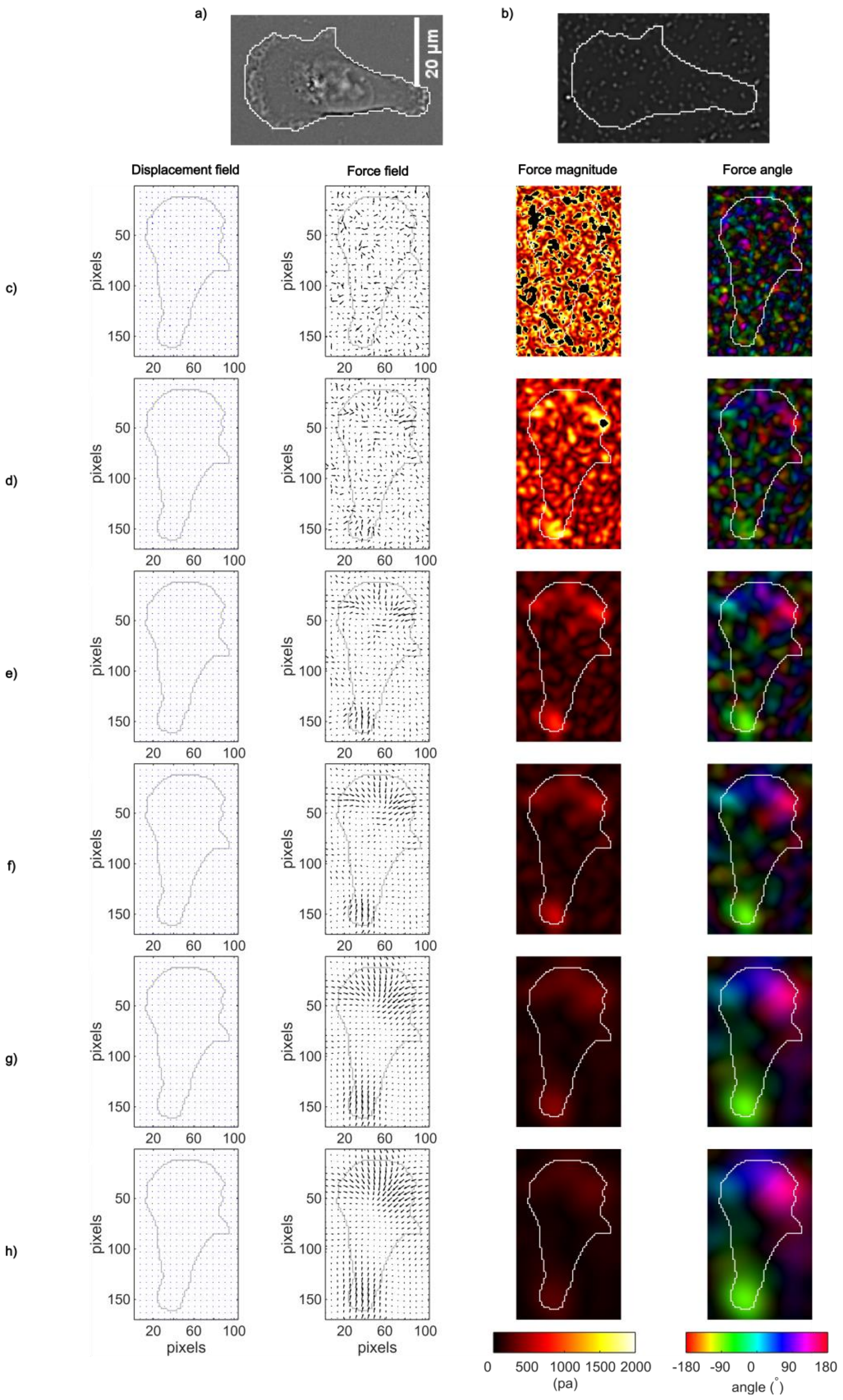


Figure S20: Effect of demons smoothing on force reconstruction for a weakly pulling MDA-MB-231 cell on a stiff substrate (46.7 kPa).

(a) Brightfield image of a MDA-MB-231 cell with segmentation overlay.
(b) Fluorescent bead images before (magenta) and after (green) traction, both processed with a Laplacian of Gaussian filter. Cell segmentation is shown in white.
(c–h) Displacement fields, reconstructed force fields (adjusted size of arrows for representation), force magnitudes, and force angles for various accumulated smoothing sizes: (c) 0.75, (d) 2, (e) 4, (f) 6, (g) 10, (h) 12.
